# Arousal as a universal embedding for spatiotemporal brain dynamics

**DOI:** 10.1101/2023.11.06.565918

**Authors:** Ryan V. Raut, Zachary P. Rosenthal, Xiaodan Wang, Hanyang Miao, Zhanqi Zhang, Jin-Moo Lee, Marcus E. Raichle, Adam Q. Bauer, Steven L. Brunton, Bingni W. Brunton, J. Nathan Kutz

## Abstract

Neural activity in awake organisms shows widespread and spatiotemporally diverse correlations with behavioral and physiological measurements. We propose that this covariation reflects in part the dynamics of a unified, multidimensional arousal-related process that regulates brain-wide physiology on the timescale of seconds. By framing this interpretation within dynamical systems theory, we arrive at a surprising prediction: that a single, scalar measurement of arousal (e.g., pupil diameter) should suffice to reconstruct the continuous evolution of multidimensional, spatiotemporal measurements of large-scale brain physiology. To test this hypothesis, we perform multimodal, cortex-wide optical imaging and behavioral monitoring in awake mice. We demonstrate that spatiotemporal measurements of neuronal calcium, metabolism, and brain blood-oxygen can be accurately and parsimoniously modeled from a low-dimensional state-space reconstructed from the time history of pupil diameter. Extending this framework to behavioral and electrophysiological measurements from the Allen Brain Observatory, we demonstrate the ability to integrate diverse experimental data into a unified generative model via mappings from an intrinsic arousal manifold. Our results support the hypothesis that spontaneous, spatially structured fluctuations in brain-wide physiology—widely interpreted to reflect regionally-specific neural communication—are in large part reflections of an arousal-related process. This enriched view of arousal dynamics has broad implications for interpreting observations of brain, body, and behavior as measured across modalities, contexts, and scales.

## Introduction

The past decade has seen a proliferation of research into the organizing principles, physiology, and function of ongoing brain activity and brain “states” as observed across various recording modalities, spatiotemporal scales, and species [1–4]. Ultimately, it is of interest to understand how such wide-ranging phenomena are coordinated within a functioning brain. However, as different domains of neuroscience have evolved around unique subsets of observables, the integration of this research into a unified framework remains an outstanding challenge [5, 6].

Interpreting ongoing neural activity is further complicated by its widespread, spatiotemporally heterogeneous relationships with behavioral and physiological variables [7–12]. For instance, in awake mice, individual cells and brain regions show reliable temporal offsets and multiphasic patterns in relation to spontaneous whisker movements or running bouts [7, 8, 13–15]. Such findings have motivated significant attention toward decomposing and interpreting neural variance that appears intermingled with state changes and movements [16–18]. Notwithstanding efforts to statistically [7, 9, 11] or mechanistically [19–22] decompose this covariance, there is lingering mystery surrounding the endogenous processes that generate and continuously orchestrate the underlying variation in time [23].

In this work, we propose a parsimonious explanation that ties together these observations: that neural activity, organismal physiology, and behavior are jointly regulated through an intrinsic, arousal-related process that continuously evolves over a latent manifold on the timescale of seconds. This unified perspective stems from the observation that nominally arousal-related fluctuations pervade—and hence, may help to bridge—effectively all domains of (neuro-)physiology and behavior [24–31]. Most recently, ongoing arousal fluctuations—indexed by scalar quantities derived from pupillometry, locomotion, cardiorespiratory physiology, or electroencephalography (EEG) [24, 32]—have emerged as a principal statistical factor of global neural variance [7, 33]. Nonetheless, the interpretation and integration of these findings is hindered by the longstanding challenge of precisely defining what “arousal” actually is [26, 34]. Although reducing this construct to individual biological mechanisms can enhance clarity, it fails to address the organism-wide coordination implicit in arousal [24]. As an alternative approach to demystify this construct, we strengthen and extend the above findings by linking variation in common arousal indices to the dynamics of a specific underlying process. We show that this process—which we refer to as *arousal dynamics*—is inherently multidimensional, and is nonlinearly coupled to observables, ultimately accounting for even greater spatiotemporal heterogeneity in brain measurements than is conventionally attributed to arousal (e.g., based on a single factor in linear regression). Crucially, we ground our explanatory framework in (data-driven representations of) this process and its couplings, rather than a statistical factor or inferred biological mechanisms.

A key motivation for our framework is to provide a unified basis for brain state research pursued across two largely separate paradigms. The first, primarily involving invasive recordings in awake mice, has established the regional dependence of cortical activity (especially at frequencies above 1 Hz) upon ongoing fluctuations in arousal and behavioral state [8, 9, 11, 35]; however, general principles have remained elusive [1, 5]. The second, originating in non-invasive human neuroimaging, has established general principles of spontaneous fluctuations in large-scale brain activity (particularly below 0.1 Hz) [36–38], which are characterized by a topographically organized covariance structure. This structure is widely thought to emerge from regionally specific neuronal interactions [39–41]; as such, arousal is not conventionally understood as fundamental to the physiology or phenomenology of interest. Instead, in this paradigm, arousal enters as either a slow or intermittent modulator of interregional covariance [31, 42–47], or as a distinct (i.e., physiologically and/or linearly separable) global component [48–51]. Yet, growing evidence indicates that global spatiotemporal processes (1) predominate at frequencies below 0.1 Hz, (2) can account for topographic structure [52–57], (3) covary with arousal indices on the timescale of seconds [37, 42, 46, 51, 54, 58], and, crucially, (4) remain dynamically interdependent, regardless of how they are decomposed [56, 59–62]. Together, we believe these themes invite a radical reinterpretation of this paradigm [63]—in particular, one which ultimately aligns this paradigm with the first. In short, we propose that (1) the spatial structure of arousal modulation arises from a complex, nondecomposable spatiotemporal process, and that (2) this process constitutes the primary generative mechanism underlying topographically organized covariance structure.

In this study, we test a central prediction of this unified account: that a scalar index of arousal suffices to accurately reconstruct multidimensional measurements of large-scale spatiotemporal brain dynamics. Inherent in this prediction is the hypothesis that “arousal”, as typically operationalized, is underpinned by a specific process that is sufficiently complex—namely, multidimensional [64]—to sustain nonequilibrium behavior, such as persistent fluctuations and traveling waves. To recover the latent dimensions of this hypothesized process, we employ a time delay embedding [65–67]—a powerful technique in data-driven dynamical systems [68] which, under certain conditions, theoretically enables reconstruction of dynamics on a manifold from incomplete, even scalar, measurements. This theory is further strengthened by the Koopman perspective of modern dynamical systems [69], which shows that time delay coordinates provide universal embeddings across a range of observables [70].

Through this framework, we provide experimental support for our predictions across simultaneous wide-field recordings of neuronal calcium, metabolism, and hemodynamics, showing that the spatiotemporal evolution of these measurements is largely reducible to a common latent process that is continuously tracked by pupil diameter. In particular, we show that a time delay embedding of pupil diameter may be used to reconstruct a latent arousal manifold; time-invariant mappings from this manifold predict substantial variation in diverse modalities both within and between mice, suggesting the arousal state-space as a generalizable embedding for brain and organismal dynamics. These findings reveal a deep connection between the mechanisms supporting brain-wide spatiotemporal organization and organism-wide regulation.

## Results

We performed simultaneous multimodal widefield optical imaging [71] and face videography in awake, head-fixed mice. Fluorescence across the dorsal cortex was measured in 10-minute sessions from a total of *N* = 7 transgenic mice expressing the red-shifted calcium indicator jRGECO1a primarily in excitatory neurons.

We observed prominent, spontaneous fluctuations in pupil diameter and whisker motion throughout each recording session (Fig. 1B). Consistent with prior work [7, 72], these fluctuations were closely tracked by the leading linear dimension of the neural data, which we obtained via traditional principal component analysis (PCA) of the widefield calcium time series (Fig. 1B). Notably, this leading component dominated the calcium time series, accounting for between 70–90% of the variance in all seven mice (Fig. S1).

**Figure 1:**
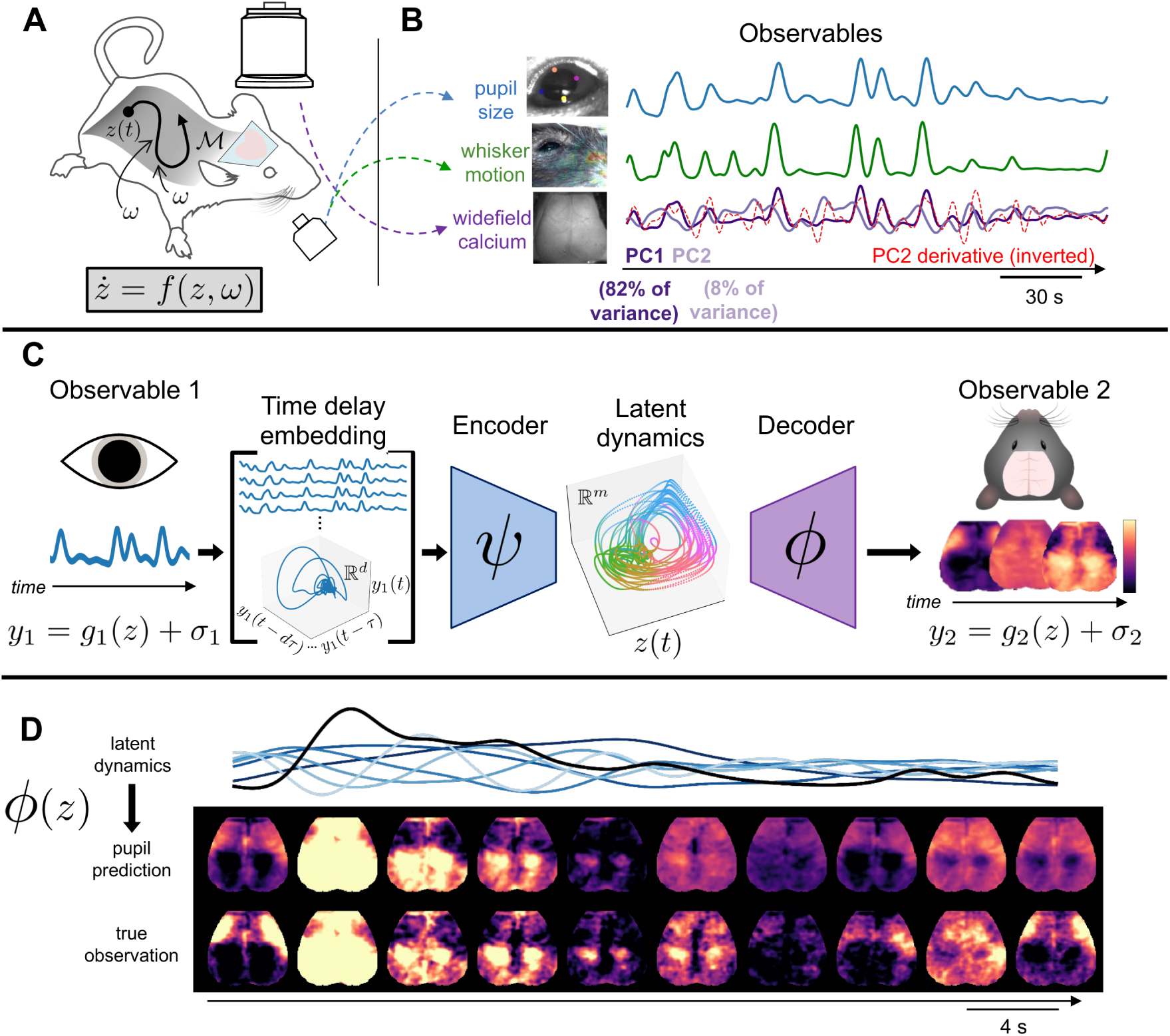
Study overview. **A** We performed simultaneous widefield optical imaging and face videography in awake mice. We interpret these measurements as observations of dynamics on an organism-intrinsic arousal manifold M (i.e., a low-dimensional surface within the state-space). Trajectories of the state *z* along M result from intrinsic arousal dynamics and various stochastic (i.e., arousal-independent) perturbations *ω*(*t*) (arising from within and outside the organism), such that 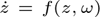. **B** Three observable quantities derived from these recordings: scalar indices of pupil diameter and of whisker motion, and high-dimensional images of widefield calcium fluorescence. Here, widefield images are represented as the time series of the first two principal components (PCs). The first brain PC (PC1) closely tracked pupil diameter and whisker motion and accounted for 70–90% of the variance in each mouse (Fig. S1). However, remaining PCs often retained a clear temporal relationship with arousal. Specifically, here PC1 closely resembles PC2’s derivative (red dashed line, Pearson *r* = −.73), implying that (nonlinear) correlation with arousal dynamics extends even beyond the dominant “arousal component” [7, 72]. **C** Our study describes an approach to parsimoniously model “arousal dynamics” and its manifestations. Each observable *yi* is interpreted as the sum of an observation function on *z* and observation-specific variability *σi* (i.e., *yi* = *gi*(*z*) + *σi*). We test the specific hypothesis that pupil size and large-scale brain activity are both regulated by a common arousal-related process. Takens’ embedding theorem [65] implies the possibility of relating these observables via a composition of encoder and decoder functions (i.e., *ψ* and *ϕ*) that map to and from a low-dimensional latent space representing the full state dynamics (*z*(*t*)). The encoder maps to this *m*-dimensional latent space from a *d*−dimensional “time delay embedding” (*d > m*) comprising *d* time-lagged copies of *y*1 (i.e., pupil diameter). **D** An example epoch contrasting original and pupil-reconstructed widefield images (held-out test data from one mouse). Black is pupil diameter; blue traces are delay coordinates, where increasing lightness indicates increasing order (Methods).

Such a dominant explanatory factor for global covariance has been widely interpreted to reflect “arousal”— a theoretical construct that, while rarely defined explicitly [34], is conventionally understood as a uni-dimensional continuum indexing the overall “wakefulness” or “activation” of an organism [24, 32, 73]. Although intuitively appealing, this framing tacitly promotes the view that brain activity naturally decomposes into linearly independent factors. In turn, this compels ad hoc explanations for the remaining orthogonal dimensions of neural data—dimensions that are, by definition, “not arousal”. Here, these dimensions correspond to the remaining PCs (e.g., [7, 9, 10]); however, orthogonality also plays a central role in defining the scope of “arousal” in psychology via factor analysis (e.g., [74]), and in the common practice of linearly regressing out arousal indices.

An important limitation of these approaches is that linear dimensions need not correspond to physically meaningful variables. More fundamentally, a linear framework imposes a notion of independence that— while mathematically convenient—is essentially arbitrary from a biological perspective, and does not imply physical or mechanistic separability [75, 76] (see also [77, Ch. 6]).

Consistent with these points, we observed that higher PCs typically retained a clear temporal relationship to PC1, often closely approximating its successive time derivatives (Fig. 1B). These dynamical relationships are consistent with our alternative view: that scalar indices of “arousal”, such as pupil size or a single PC, are best understood as one-dimensional projections of an underlying process—one whose full manifestations extend across multiple (Euclidean) dimensions (Fig. 1A). Crucially, on this view, the corresponding dimensions of the data remain fundamentally entangled, registering different aspects of a common process.

We refer to this underlying process as “arousal dynamics”. Our core hypothesis is that the infra-slow spatiotemporal evolution of large-scale brain activity is primarily a reflection of this multidimensional, unified process. Below, we motivate this idea from a physiological perspective before introducing its dynamical systems framing. Ultimately, this framing enables us to empirically evaluate predictions motivated by this perspective.

### Studying arousal dynamics

We interpret nominally arousal-related observables (e.g., pupil size, whisking) to reflect in large part the dynamics of a unified, organism-wide regulatory process (Fig. 1A). This dynamic regulation [78] may be viewed as a low-dimensional process whose evolution is tied to stereotyped variations in the high-dimensional state of the organism. Physiologically, this abstraction captures the familiar ability of arousal shifts to elicit multifaceted, coordinated changes throughout the organism (e.g., “fight-or-flight” vs. “rest-and-digest” modes) [79]. Operationally, this framing motivates a separation between the effective dynamics of this system, which can be represented on a latent, low-dimensional manifold [80–82], and the high-dimensional “observation” space, which we express via time-invariant mappings from the latent manifold. In this way, rather than assuming high-dimensional, pairwise interactions among the measured variables, we attempt to parsimoniously attribute widespread variance to a common, latent dynamical system [83, 84].

To specify the instantaneous latent arousal state, our framework takes inspiration from Takens’ embedding theorem [65], which provides general conditions under which the embedding of a scalar time series in time delay coordinates preserves the topological properties of (i.e., remains diffeomorphic to) the attractor in the full state-space of the original system. Delay embedding has become an increasingly powerful technique in modern data-driven dynamical systems [68]. Specifically, time delay coordinates have been shown to provide universal coordinates that represent complex dynamics for a wide array of observables [70], related to Koopman operator theory in dynamical systems [69, 85].

As we hypothesize that brain-wide physiology is spatiotemporally regulated in accordance with these arousal dynamics [63], Takens’ theorem leads us to a surprising empirical prediction: a scalar observable of arousal dynamics (e.g., pupil diameter) can suffice to reconstruct the states of a high-dimensional observable of brain physiology (e.g., optical images of cortex-wide activity), to the degree that the latter is in fact coupled to the same underlying dynamical process [67, 86, 87].

To test this hypothesis, we developed a framework for relating observables that evolve in time according to a shared dynamical system. Fig. 1C illustrates our data-driven approach to compute these relationships. In each mouse, we embed pupil diameter measurements in a high-dimensional space defined by its past values (i.e., time delay coordinates). We wish to discover a map (or encoder) *ψ* from this newly constructed observation space to a low-dimensional latent space *z*, which represents the instantaneous arousal state as a point within a continuous, observation-independent state-space. We additionally seek a mapping (decoder) *ϕ* from this latent space to the observation space of widefield calcium images. These mappings can be approximated with data-driven models learned using linear or nonlinear methods. Ultimately, this procedure enables us to express each instantaneous widefield calcium image as a function of a multidimensional state defined by pupil delay coordinates, to the extent that these two observables are deterministically linked to the same latent arousal state *z*.

### Spatiotemporal dynamics from a scalar

For all analyses, we focused on (“infra-slow” [88, 89]) fluctuations *<*0.2 Hz, as this frequency range distinguishes arousal-related dynamics from physiologically distinct processes manifesting at higher frequencies (e.g., delta waves) [53, 90]. To model the hypothesized mappings, we implemented an iterative, leave-one-out strategy in which candidate models were trained using 10-minute recordings concatenated across six of the seven mice, and subsequently tested on the held-out 10-minute recording obtained from the seventh mouse. This modeling pipeline formed the basis for all main text results. We also implemented a separate, within-mouse pipeline in which we trained models on the first 5 minutes of data for each mouse and report model performance on the final 3.5 minutes of the 10-minute session. Results from this within-mouse pipeline are included in the Supplementary Appendix (Fig. S9). Details for both modeling pipelines are provided in the Methods.

For both pipelines, candidate models included a linear regression model based upon a single lag-optimized copy of pupil diameter (“No embedding”), as well as a model incorporating multiple delay coordinates and nonlinear mappings (“Latent model”). Importantly, both models allow us to assess the hypothesis that the widefield measurements primarily reflect arousal-related variation. However, the latter model adds the ability to capture complex aspects of pupil-widefield covariation that are more uniquely predicted by our dynamical view of arousal, enabling stronger validation of this particular interpretation. Nonlinear mappings were learned through simple feedforward neural networks, which were trained within an architecture modeled after the variational autoencoder [91] (with hyperparameters fixed across mice; Methods). See Supplementary Appendix (Supplementary Text and Figs. S2-S3) for validation of this framework on a toy (stochastic Lorenz) system.

Modeling results from the leave-one-out pipeline are shown in Fig. 2. We first considered the overall variance explained, summarized across time points and pixels. In each of *N* = 7 mice, we succeeded in reconstructing the majority of the variance in widefield calcium images on the basis of (simultaneously measured) pupil diameter, accounting for 60–85% of the total variance *<*0.2 Hz (Fig. 2A). As noted above, the total variance was dominated the first PC (Fig. S1), which showed a tight, linear relationship with pupil diameter. Accordingly, much of the total variance could also be modeled via linear regression with a one-dimensional pupil regressor (Fig. 2A). Thus, heuristically, both models support the notion that widefield calcium measurements are dominated by nominally “arousal-related” variation. However, as the total variance effectively treats each time point and pixel independently, this quantity fails to capture the dynamical and geometric structure inherent in the data—properties that may more directly relate to the underlying, spatiotemporally embedded generative process. Therefore, we next examined how well each model preserved these relational aspects of the data.

**Figure 2:**
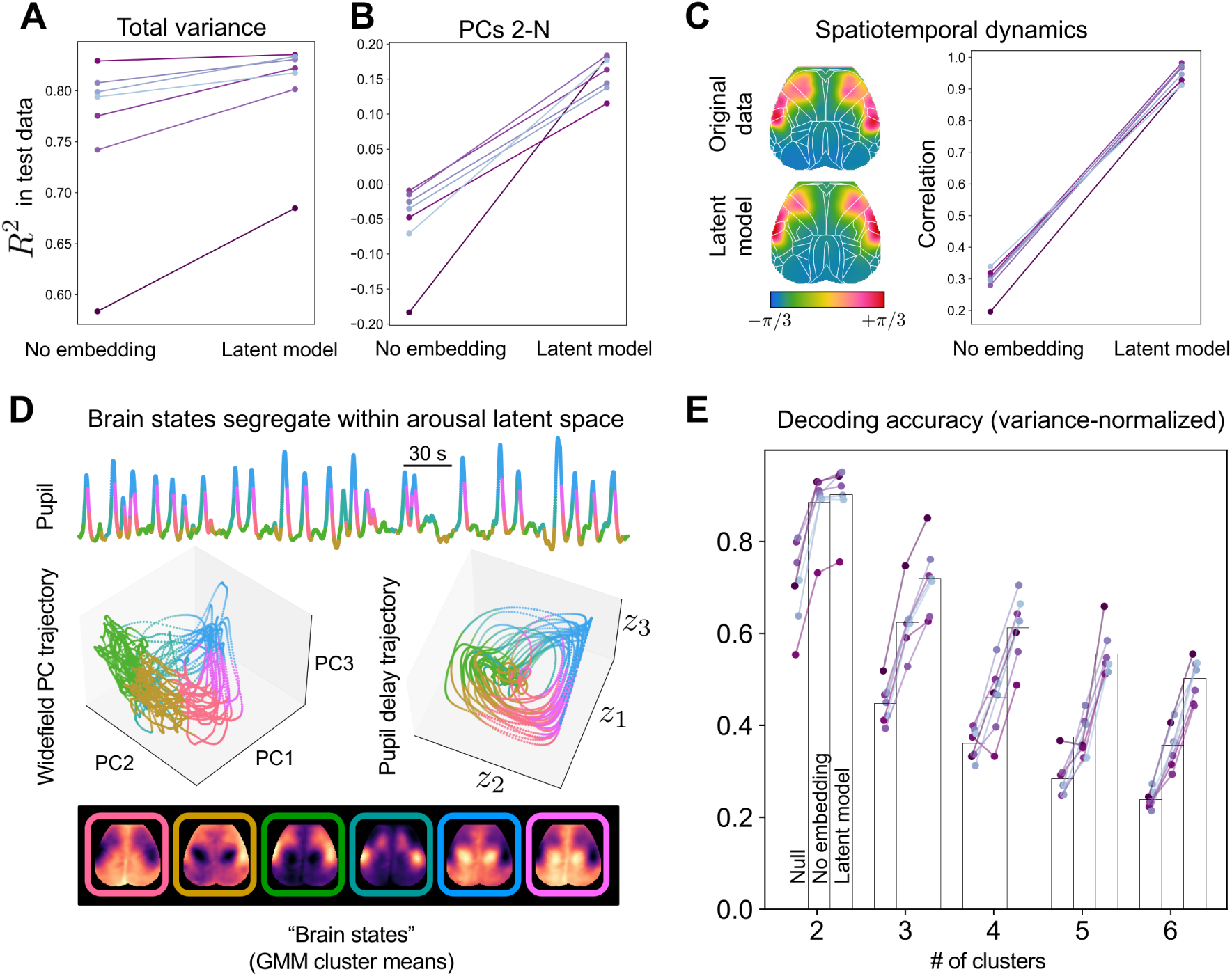
Pupil diameter enables reconstruction of spatiotemporal brain dynamics. **A** Variance explained in held-out widefield data from each mouse (for each prediction, the model was trained on data from the other 6 mice). A simple linear regression model based on one (lag-optimized) pupil regressor (“No embedding”) accurately predicted ∼ 70–85% of variance in six of seven mice. Delay embedding and nonlinearities (“Latent model”) extended this level of performance to all mice while further augmenting model performance, primarily by accounting for variance spread along higher dimensions (PCs 2-N) (B). See Fig. S6 for pixelwise maps of variance explained. **C** Analysis of propagation dynamics. Phase maps were obtained from dynamic mode decomposition applied to the original and reconstructed data, and subsequently compared via (circular) spatial correlation. Phase maps from one example mouse are shown on the left (phase maps from the 1D model are not shown, as they lack phase structure due to the absence of complex-valued modes). Allen Mouse Common Coordinate Framework atlas overlaid in white. **D** Example state trajectories (held-out data) visualized in coordinates defined by pupil diameter (top), the first three widefield PCs (left) or three latent variables obtained from the delay-embedded pupil time series (right). Trajectories are color-coded by cluster identity as determined by a Gaussian mixture model (GMM) applied to widefield image frames (*k* = 6 clusters). Cluster means are shown below these trajectory plots. Color segregation in pupil delay coordinates indicates spatially-specific information encoded in pupil dynamics. Similar color arrangement within both embeddings suggests their shared topological properties. **E** Decoding “brain states” from pupil dynamics. Comparison of decoder accuracy in properly identifying GMM cluster assignments for each image frame based on the most frequent assignment in the training set (“Null” decoder), the “No embedding” model, or the “Latent model”.

Crucially, we found that including multiple delays and nonlinearities additionally captured substantial variance orthogonal to the first PC (akin to residuals following global signal regression) (Fig. 2B) (see also Figs. S4, S5, and S9). Spatially, the topography of total explained variance was closely related to the first PC and strongly weighted toward posteromedial cortices (Fig. S6), resembling a familiar arousal-related topography as estimated by linear mappings from arousal indices [8, 9, 11, 92, 93]. However, in contrast, delays and nonlinearities allowed the model to also explain variance in anterolateral (“orofacial”) cortices (Fig. S6), thus complementing the conventional arousal-related topography. The pattern of additional explained variance resembled the spatial correlates of pupil derivatives and pupil amplitude envelope (Fig. S7)—two features which are implicitly represented in the delay embedding, and which are known to be tracked by neuromodulator dynamics [94, 95]. Interestingly, the largest increases in explained variance were consistently observed within a spatially focal set of regions that appeared to intersect the two main topographies (i.e., posteromedial and anterolateral cortices) (Fig. S6).

These consistent patterns suggest that modeling improvements did not trivially result from increased model complexity. Indeed, shuffling pupil and widefield measurements across mice precluded any ability to learn models that generalized to held-out data (Fig. S8). We conclude that an appreciable amount of widefield variance lying outside the dominant PC retains a time-invariant relationship with pupil dynamics (Fig. 2B-C).

The ability to model variance along multiple orthogonal dimensions of the widefield measurements indicates shared multidimensional, nonlinear structure between widefield and pupil dynamics. Nonetheless, this still does not directly address whether the model predictions faithfully reproduce the bona fide *spatiotemporal dynamics* of the widefield data, as would be expected if pupil and widefield measurements indeed share a common generative dynamical mechanism that is accurately captured by our model. To test this hypothesis more directly, we approximated propagation structure by fitting a linear dynamical systems model (via dynamic mode decomposition [69]; Methods) to both the original and reconstructed widefield measurements, then examined the leading spatial mode of each fitted dynamical system. In all seven mice, the original widefield data yielded a complex-valued leading spatial mode revealing a stereotyped pattern of cortical propagation consistent with prior observations [53, 96]. Crucially, data reconstructed by the “Latent model” consistently preserved this global propagation structure, as evidenced by the spatially similar phase patterns exhibited by the leading mode from each mouse (Fig. 2C). In contrast, the “No embedding” model failed to capture spatiotemporal dynamics altogether (see also Supp. Videos 1-2). This failure is expected, as bona fide spatiotemporal dynamics are inherently multidimensional: a one-dimensional dynamical system can yield only real-valued modes (eigenvectors), whereas true spatiotemporal dynamics correspond to interregional phase shifts that exist in the complex plane. Thus, generating and representing multiple distinct phase shifts necessitates at least two (Euclidean) dimensions.

Analyses based on hidden Markov models (HMMs) further supported the unique ability of delay embeddings to preserve the dynamical structure of the original widefield measurements (Fig. S10). Collectively, these findings and theoretical insights indicate that multiple orthogonal dimensions of brain measurements are inherently coupled due to an underlying spatiotemporal process—one that is faithfully reflected in the continuous evolution of the pupil.

Having successfully accounted for the majority of the variance, along with multidimensional and dynamical structure, we next asked whether these results could be attributed to high-amplitude, temporally isolated events, or whether they reflect a continuous coupling between the spatial topography of calcium measurements and the instantaneous state of arousal dynamics. To address this question, we evaluated the correspondence between the original and reconstructed datasets on a frame-to-frame basis, giving each time point (and thus, each image frame) equal weighting. Specifically, we first normalized each image frame to unit variance and projected it onto the top three PCs, then assigned these normalized, dimensionally-reduced images to clusters by fitting a Gaussian mixture model (GMM) (Methods). In this way, image frames were assigned to clusters based purely on their spatial similarity. Following this procedure, we assessed whether pupil dynamics allowed us to decode the cluster assignment of widefield data on a frame-to-frame basis.

Fig. 2D-E illustrates the results of these clustering and decoding analyses. Fig. 2D demonstrates the segregation of distinct spatial topographies to different regions of the arousal latent space (derived from pupil diameter), indicating that arousal dynamics are informative of spatially-specific patterns of cortex-wide activity. Fig. 2E shows decoding accuracy in test data as a function of the number of allowable clusters. Decoding accuracy was computed for each model as the percentage of “ground truth” GMM cluster assignments that matched the predictions generated by the same GMM conditioned on the reconstructed (rather than the original) image frames. A “null” score was obtained by assigning all frames to the most frequent assignment in the ground truth data.

In general, we observed that spatial patterns largely alternated between two primary patterns, consistent with large-scale spatiotemporal measurements reported across modalities and species [62, 63, 92, 97]. With a two-cluster solution, pupil dynamics enabled accurate assignment of more than 80% of image frames (Fig. 2E). Decoding accuracy remained high with three distinct topographies, with the latent model enabling an average accuracy of *>*70%. With more than three clusters, we found spatial topographies to be increasingly less distinctive, leading to gradually worsening decoding accuracy. Yet, decoding accuracy continued to follow a common trend, steadily improving from the “Null” decoder to the “No embedding” decoder, and, finally, to the “Latent model” decoder. The superior accuracy of the latter was primarily rooted in its ability to simultaneously track spatial variation along multiple orthogonal dimensions, rather than just the first PC. Indeed, discrepancies in prediction accuracy became even more pronounced when assigning equal weight to the three PCs used for the clustering analysis (Fig. S11). In addition, we observed similar or stronger decoding advantages of the latent model when labeling states according to an HMM rather than a GMM (Fig. S10). These results confirm that pupil dynamics reliably index the instantaneous spatial topography of large-scale calcium fluorescence.

Collectively, our analyses support the interpretation that arousal dynamics, as indexed by pupil diameter, represent the predominant physiological process underlying cortex-wide spatiotemporal dynamics on the timescale of seconds.

### Multimodal spatiotemporal dynamics from a shared latent space

We have hypothesized that arousal dynamics support the spatiotemporal regulation of brain-wide *physiology* —thus extending beyond the activity of neurons per se (e.g., [29, 72, 98]). Spatially structured fluctuations in metabolic and hemodynamic measurements are also well-established, and are generally interpreted to reflect state-, region-, and context-dependent responses to local neuronal signaling [99]. On the other hand, our account posits that multiple aspects of physiology are jointly regulated according to a common process. This suggests the possibility of more parsimoniously relating multiple spatiotemporal readouts of brain physiology through a common low-dimensional latent space reconstructed from pupil diameter.

To test this prediction, we took advantage of the additional readouts provided by the multispectral optical imaging platform [71]—namely, oxidative metabolism (as indexed by flavoprotein autofluorescence, or FAF [100–102]) and hemodynamics (which approximates the physiological basis for fMRI), which were obtained simultaneously with the neuronal calcium measurements. We attempted to reconstruct all three observables on the basis of delay embedded pupil measurements. Importantly, we did not retrain the encoder *ψ* learned in the context of predicting calcium measurements in Fig. 2. Rather, we simply trained additional decoders to map from the latent space defined by *ψ* to spatiotemporal measurements of FAF and oxygenated hemoglobin (*ϕ*_2_ and *ϕ*_3_ in Fig. 3A, respectively).

**Figure 3:**
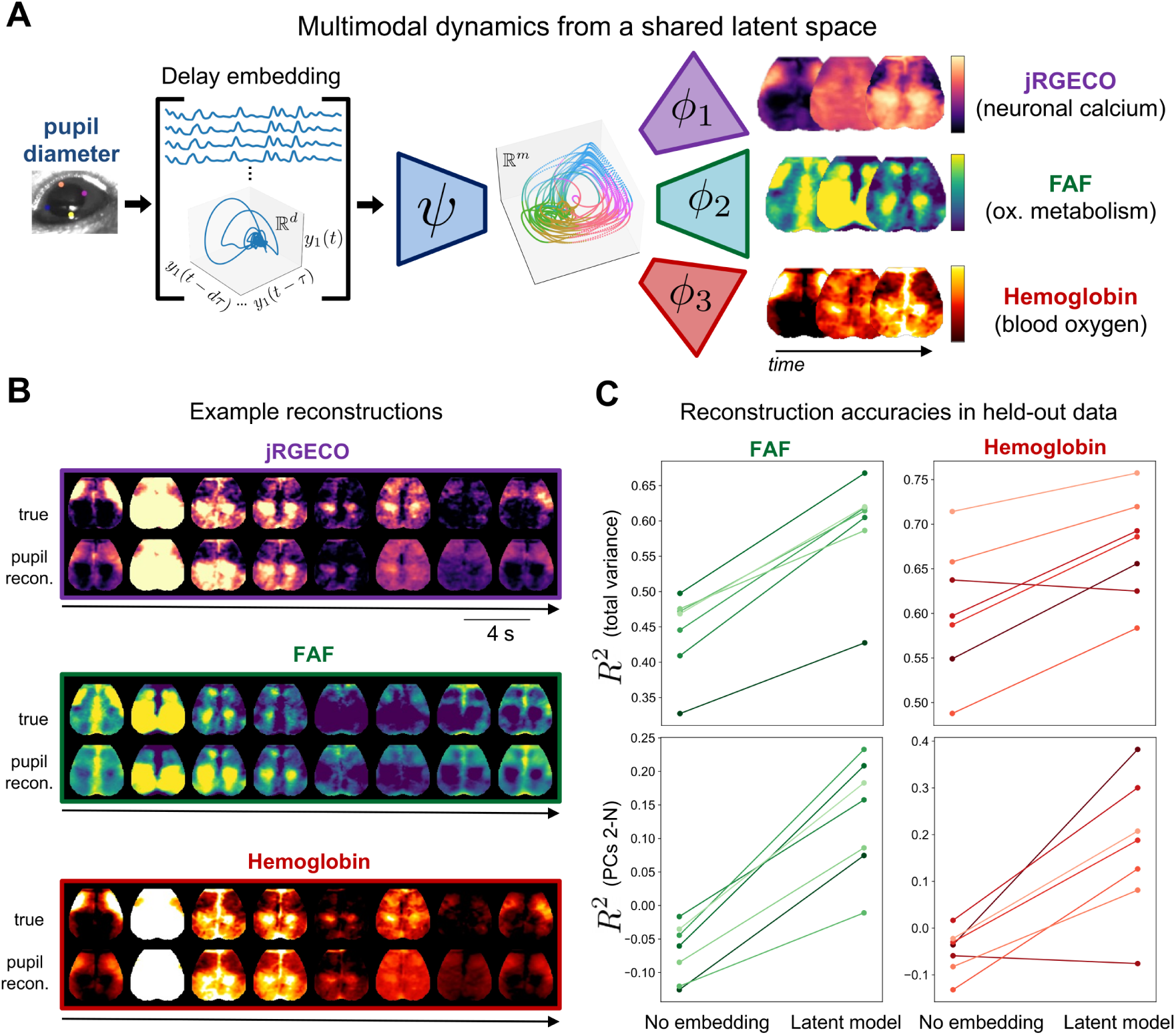
Multimodal physiological measurements can be reconstructed from a shared latent manifold. **A.** The delay embedding framework extended to multimodal observables. We used the same encoder previously trained to predict calcium images. We coupled this encoder to decoders trained separately to predict FAF and hemoglobin. **B.** Example reconstructions of neuronal calcium (jRGECO fluorescence), metabolism (FAF), and blood oxygen (concentration of oxygenated hemoglobin). **C.** Reconstruction performance for each observable (as in Fig. 2A). See Fig. 2 for results with calcium, and Fig. S14 for pixelwise maps of variance explained.

Modeling accuracy for FAF and hemoglobin measurements are shown in Fig. 3B. As with calcium fluorescence, we were able to accurately model the large-scale spatiotemporal dynamics of these two measurements in each of *N* = 7 mice. These measurements were similarly dominated by the first PC (Figs. S12 & S13). Interestingly, however, for these measurements, delay embeddings and nonlinearities made a much larger contribution to the explained variance, even after adjusting for a globally-optimized delay (Figs. 3C & S14). Thus, a far larger portion of metabolic and hemodynamic fluctuations can be attributed to a common, arousal-related mechanism than is conventionally recognized (see also [54, 63]). For context, predictions based on pixelwise calcium signals are shown in Fig. S15). We revisit the physiological interpretation of these relationships in the Discussion.

### Unifying observables through arousal dynamics

Analyses thus far support our dynamical systems interpretation of arousal, which offers a parsimonious account of ongoing fluctuations in multimodal measurements of large-scale spatiotemporal brain activity. A key motivation for this framing is the growing diversity of observables linked to arousal indices [6], despite the persistent lack of clarity surrounding this construct [34]. Accordingly, we now consider how our framework can quantitatively integrate such findings via a unified representation of arousal.

Crucially, by relating observables through an organism-intrinsic manifold, rather than an imposed stimulus or task (cf. [103]), our framework has the potential to generalize across different experimental datasets, enabling integration of our findings into a more holistic context than is feasible in any single experimental study. To explore this potential, we extended our framework to publicly available behavioral and electrophysiological recordings from the Allen Institute Brain Observatory [104]. This dataset includes 30-minute recording sessions of mice in a task-free context, i.e., without explicit experimental intervention. From this dataset, we computed eight commonly used indices of brain state and arousal (see Methods): pupil diameter, running speed, hippocampal *θ/δ* ratio, the instantaneous rate of hippocampal sharp-wave ripples (SWRs), bandlimited power (BLP) derived from the local field potential (LFP) across visual cortical regions, and mean firing rate across several hundred neurons per mouse recorded with Neuropixels probes [105]. The two hippocampal indices were both derived from the LFP of the CA1 region. BLP across visual cortex was further analyzed within three canonical frequency ranges: 0.5–4 Hz (“delta”), 3–6 Hz (“alpha” [106]), and 40–100 Hz (“gamma”).

We began by concatenating measurements across mice, then trained neural networks to predict each observable from a latent space constructed from (delay embedded) pupil measurements. After training on concatenated measurements, we applied the learned encoding to the delay embedded pupil measurements of each individual mouse, training decoders to map from this shared latent space to the mouse’s own observables. Finally, we repeated this procedure for the primary widefield dataset—thus using pupil diameter as a common reference to align observables across datasets (and mice) to a shared manifold (see Methods for further details).

Fig. 4 illustrates the resulting, multi-dataset generative model. For visualization, we defined the arousal manifold M as a 2D latent space parameterized by two discovered coordinates, *z*_1_ and *z*_2_. Next, we characterized M in terms of each observable *y_i_* by evaluating the corresponding mappings *ϕ_i_* at each point within a 2D grid of sample points within the latent space. Finally, we obtained a data-driven analytic representation of the latent dynamics [107] (Methods), allowing us to compute the vector field *f* (*z*) at each point within the 2D grid. Taken together, at each point along M, Fig. 4 depicts the conditional expectation for (1) the direction of flow of the latent arousal dynamics (via *f*), and (2) the state of each observable *y_i_* (via *ϕ_i_*). In this way, M, *f*, and the mappings 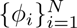 together encapsulate our dynamical, relational interpretation of arousal.

**Figure 4:**
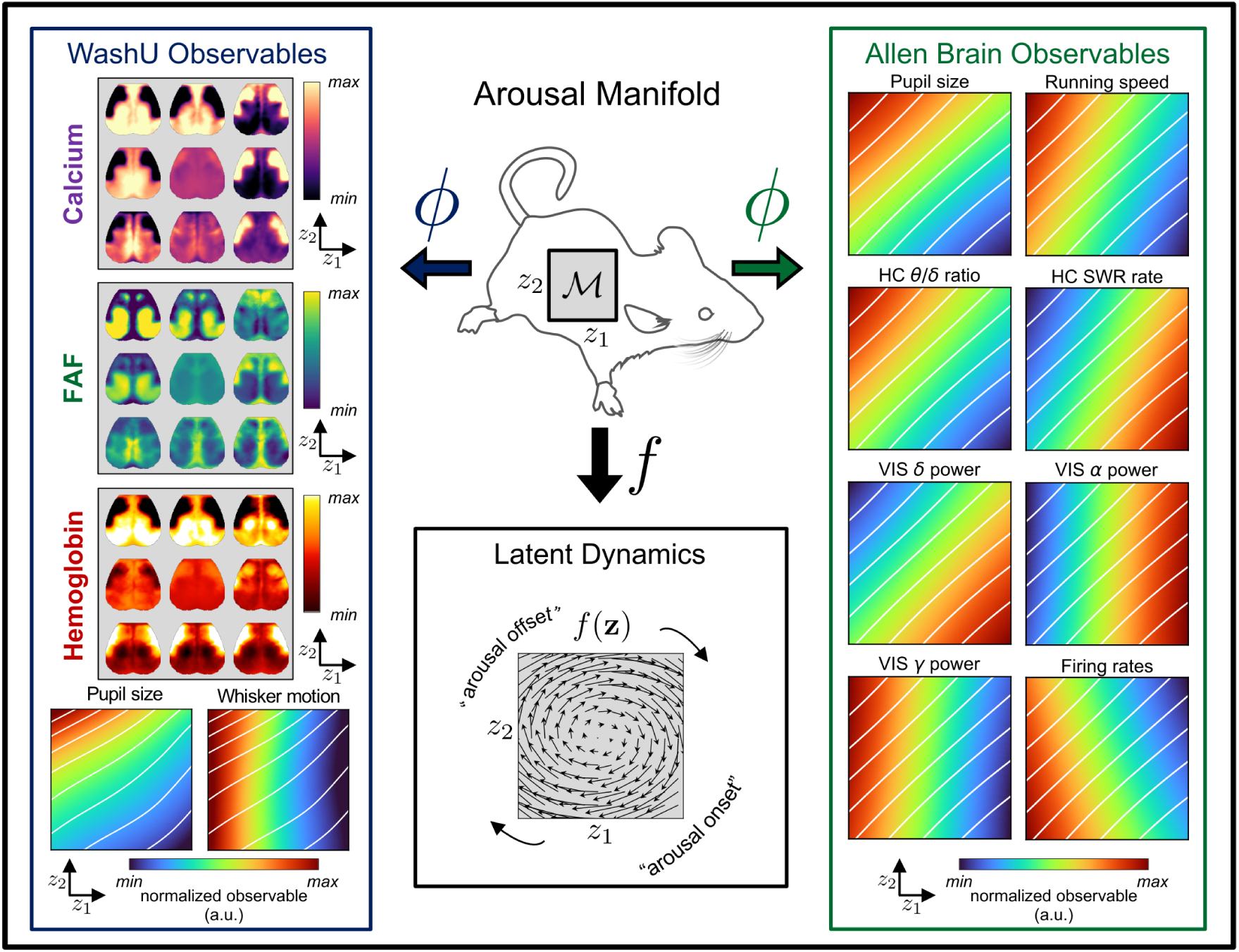
Unifying observables through arousal dynamics. The arousal manifold provides an organism-intrinsic reference frame to align and aggregate measurements across datasets. To visualize the discovered relations, we begin by defining the arousal manifold as a 2D latent space parameterized by the coordinates *z*1 and *z*2 (i.e., **z** ∈ M ⊂ R^2^) (upper middle). Each point **z** in this 2D manifold represents a latent arousal state. The learned decoders *ϕ* map each of these points to an expected value of a particular observable (e.g., a single widefield calcium image, or a single value of pupil size), thus enabling representations of the 2D arousal manifold from the perspective of various observables (left and right columns). For visualization, the widefield observables are evaluated at 3 × 3 evenly spaced points within the 2D latent space, whereas each scalar observable is evaluated over a grid of 100 × 100 sample points). White diagonals indicate iso-contours of pupil size in either the WashU or Allen Brain Observatory datasets, allowing a common visual reference across observables within the same dataset. Meanwhile, *f* (**z**), which we approximate via a data-driven analytic representation [107], maps each point in the latent manifold to an expected direction of movement (i.e., 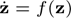), enabling the manifold to be represented also as a vector field (lower middle). Together, these representations depict how a typical sequence of arousal states, as determined by the latent, intrinsic arousal dynamics, would manifest in terms of changes to observable quantities. For instance, beginning at “arousal onset” (lower right corner of each square in the figure), the clockwise rotation indicated by arrows in *f* (**z**) is expected to manifest as an increase and subsequent decrease in whisker motion; similar changes are expected in pupil size, but slightly delayed, as the max pupil size is rotated clockwise from max whisker motion. FAF, flavoprotein autofluorescence; HC, hippocampus; SWR, sharp-wave ripple; VIS, visual cortex.

The resulting representation unifies and clarifies relations among several themes appearing in different sectors of the experimental literature, which we briefly summarize here. As expected, running speed, hippocampal *θ/δ* ratio, cortical gamma BLP, and mean firing rate are all positively correlated with pupil diameter along a single, principal dimension, *z*_1_ (i.e., the horizontal coordinate in each plot) (Fig. 4, “Allen Brain Observables”). In contrast, delta BLP, alpha BLP, and hippocampal SWR rate—all conventionally associated with internally-oriented functionalities and momentary sensory disengagement—are inversely correlated with pupil diameter along the same *z*_1_ dimension [13, 108, 109].

A second dimension (*z*_2_, vertical coordinate in each plot) clarifies more subtle temporal relationships, revealing a canonical cycle that captures diverse phenomenology previously reported in the context of brain and behavioral state transitions [94, 95, 110, 111]. Thus, the “arousal onset” (Fig. 4) begins with decreasing alpha BLP [111] and subsequent increases in firing rates and gamma BLP, which precede an increase in pupil diameter [72, 110]. The “WashU observables” indicate that this arousal onset period is also characterized by the onset of whisking and a propagation of cortical activity toward posterior regions [19, 53, 92], with calcium changes briefly lagged by metabolic and then hemodynamic changes.

Continuing clockwise from this “arousal onset” period, the Allen Brain Observables indicate roughly synchronous increases in pupil diameter, locomotion, and hippocampal *θ/δ* ratio. After these observables reach their maximum, alpha BLP begins to increase, consistent with the appearance of this cortical oscillation at the offset of locomotion [106] (marked as “arousal offset” in Fig. 4). This is succeeded by a general increase in delta BLP and hippocampal SWR rate. Across the cortex, this behavioral and electrophysiological sequence at “arousal offset” (as indicated by the Allen Brain Observables) is associated with a propagation of activity from posterior to anterior cortical regions (as indicated by the WashU observables). Finally, activity appears across lateral orofacial cortices, which remain preferentially active until the next arousal onset. This embedding of cortex-wide topographic maps within a canonical “arousal cycle” resembles prior results obtained from multimodal measurements in humans and monkeys, which additionally link these cortical dynamics to topographically parallel rotating waves in subcortical structures [63].

Taken together, this framework provides a unified, parsimonious, and quantitative representation of a functionally coherent spatiotemporal pattern of changes that is registered only piecemeal through diverse experimental modalities. Although verifying this model would ideally require simultaneous measurements of all observables—which remains practically infeasible—the ability to integrate partial observations from different experiments plays a key role in contextualizing disparate observations, facilitating hypothesis generation, and enabling large-scale model construction, thereby laying groundwork for future experiment-theory iteration.

## Discussion

We have proposed to interpret arousal as a specific, multidimensional process that spatiotemporally regulates brain-wide physiology in parallel with organismal physiology and behavior. We support this account through a data-driven modeling framework, demonstrating that a scalar index of arousal, pupil diameter, suffices to reconstruct the multidimensional state dynamics underlying the spatiotemporal evolution of cortex-wide physiology. We interpret these results as support for the hypothesis that spontaneous, infraslow fluctuations in large-scale brain activity primarily reflect an intrinsic regulatory mechanism [63]. Collectively, these insights carry important implications for understanding brain states and arousal, large-scale brain dynamics, and the interrelations among multimodal readouts of brain physiology. These implications are considered in what follows.

The concept of “brain state” has gained heightened significance in systems neuroscience as experimental advances have made studies in awake, behaving organisms increasingly routine [5, 6, 35]. Although brain state is typically viewed as comprising several biological, cognitive, or behavioral factors—one of which is often presumed to be “arousal” [1, 10, 35]—it has remained unclear how to reliably decompose brain state without more precisely defined factors. Construing organisms as dynamical systems [83, 112], our approach has been to factorize brain (or rather, organismal) state in terms of constituent endogenous processes—one of which, examined herein, we identify with the arousal construct. Despite grounding arousal in a specific process, our results reveal this process to be multidimensional and nonlinearly coupled to various observables—ultimately implying even broader relevance for arousal than presently recognized.

Our generalized view of arousal resonates with two emerging themes in the literature. First, brain or arousal states appear to represent distinct (neuro-)physiological regimes, such that state changes are associated with sweeping changes in the behavior of faster timescale processes [29, 113, 114]. This limits the ability to account for arousal-dependence by simply orthogonalizing observations with respect to a scalar arousal index (e.g., via linear regression of pupil diameter or removal of the leading PC; Fig. 2B). Second, numerous studies have demonstrated non-random transitions among brain states [1, 12, 115, 116], along with the functional relevance of *dynamical* indices of arousal, such as the phase or derivative of pupil size [72, 94, 110, 117], often with surprising timescale-invariance [14, 63, 90]. These observations imply an underlying, nonlinear dynamic process [118]. Together, these two themes suggest a view of arousal as constituting a complex, pervasive, and persistent dynamic “background”; this complicates the ability to interpret spontaneous or task-related [9, 11] variation observed over similar timescales—that is, even on the order of seconds—without reference to this changing *internal context* [2, 17]. To this end, modeling approaches such as in Fig. 4 can refine interpretations by more rigorously conditioning observations upon this nonlinear dynamic background.

Indeed, by more rigorously capturing the scope of arousal dynamics, our models reveal surprisingly rich spatiotemporal structure across the cortex, prompting reconsideration of the mechanisms typically assumed responsible. Thus, over decades of research spanning multiple experimental communities, large-scale spatiotemporal brain dynamics have been overwhelmingly viewed as summations of diverse interregional interactions [37, 39, 41, 119], fueling a proliferation of decomposition techniques. Although distinctions among modalities and species should not be overlooked, evidence across these domains has increasingly converged toward a sparse set of core underlying spatiotemporal components [52, 56, 62], alongside growing appreciation for arousal-related contributions [31, 43, 46, 51]. We extend these themes by advocating a more radically unified view: that these core spatiotemporal components are themselves projections of a nondecomposable dynamical system, and that this system effectively underpins arousal. In this sense, arousal dynamics directly account for the observed spatiotemporal structure—i.e., not (primarily) by reconfiguring interregional neuronal interactions, but by actively generating spatiotemporal structure through the synchronized [87] propagations of multiple topographically organized rotating waves [63].

Importantly, our unified model specifically concerns infra-slow spatiotemporal brain dynamics. Significantly faster processes (particularly those above 1 Hz) are effectively decoupled from this timescale and likely involve distinct physiological mechanisms [90], including the kinds of flexible and regionally specific interactions that we argue play a smaller role at infra-slow timescales. Interestingly, widefield optical recordings reveal similar spatial patterns across both timescales [120], likely reflecting shared anatomical constraints (in our model, such constraints are implicitly subsumed by the time-invariant mapping *ϕ*). Nonetheless, temporal dynamics among these spatial patterns seem to differentiate the two timescales [53, 96], supporting their distinct underlying mechanisms. Still, rather than being entirely separate, spatiotemporal dynamics above 1 Hz are strongly modulated by infra-slow (and/or arousal) dynamics [53, 120, 121]. Likewise, a rich literature describes the amplitude modulation of higher-frequency brain activity according to the phase of infra-slow fluctuations [89]. We suggest that these multiscale relationships are special cases of a more general pattern by which the infra-slow evolution of arousal state modulates the parameters of faster timescale processes throughout the brain and body. Indeed, the conventional brain state indices in Fig. 4 can be viewed as precisely such parameters (e.g., “rate” and “power”). This pattern suggests multiscale extensions of the present framework, wherein dynamical models describing faster processes can be jointly parameterized by the state of a common arousal process.

This theoretical framing also leads us to a simplified view of the interrelations between electrical, metabolic, and vascular cerebral activity. Although brain function necessitates tight coupling among all three aspects of physiology, their intersection constitutes an extraordinarily high-dimensional space of biochemical interactions. These interactions are commonly viewed in terms of complex, region-dependent, and still incompletely understood neurometabolic and neurovascular cascades operating locally throughout the brain [99], posing an enormous regulatory challenge. On the other hand, there is longstanding evidence that neuromodulatory processes can directly influence neural, metabolic, and vascular activities in parallel [122–125], suggesting mechanisms enabling their coordinated regulation. Based on the strong performance of our model (Fig. 3), we propose that such mechanisms are actively harnessed to effectively streamline cerebral regulation in accordance with arousal dynamics—for instance, through spatiotemporally coordinated changes to high-level parameters such as neuronal gain [126] and metabolic fluxes [127]. This strategy would reduce reliance on more conventionally studied feedforward responses to local neuronal activity (e.g., “the hemodynamic response”), which may come secondary to more integrative mechanisms supporting the continuous, dynamic, and globally coordinated nature of brain physiology (see also [29, 128]). This interpretation could help explain observations that appear more puzzling when viewing spontaneous hemodynamic fluctuations as mere responses to local neuronal activity (e.g., [129]). Ultimately, targeted experiments are needed to determine the extent to which neuromodulatory processes directly and indirectly contribute to neurometabolic and neurovascular coupling. Regardless, our results indicate that the observed dynamics and relationships are consistent with widespread coupling to a global regulatory process.

Finally, although we have adopted the term “arousal” given widespread adoption of this term for the observables discussed herein, our results are obtained through an entirely data-driven approach: “arousal” is simply the label we assign to the latent process inferred from the observables. In other words, we make no a priori assumptions about how arousal might be represented in these observables in comparison to other behavioral processes or states [130]. For this same reason, our results are likely relevant to a range of cognitive and behavioral processes that are traditionally distinguished from arousal (e.g., “attention”), but which may be intimately coupled (or largely redundant) with the dynamical process formalized herein [2, 131].

## Methods

### Datasets and preprocessing

#### Dataset 1: Widefield optical imaging (WashU)

##### Animal preparation

All procedures described below were approved by the Washington University Animal Studies Committee in compliance with the American Association for Accreditation of Laboratory Animal Care guidelines. Mice were raised in standard cages in a double-barrier mouse facility with a 12 h–12 h light/dark cycle and ad libitum access to food and water. Experiments used N=7 12-week old mice hemizygous for Thy1-jRGECO1a (JAX 030525) on a C57BL/6J background, enabling optical imaging of the jRGECO1a fluorescent calcium sensor protein primarily expressed in excitatory neurons of cortical layers 2/3 and 5 [132]. Prior to imaging, a cranial window was secured to the intact skull of each mouse with dental cement under isoflurane anesthesia according to previously published protocols [133].

##### Data acquisition

Widefield imaging was conducted on a dual fluorophore optical imaging system; details of this system have been presented elsewhere [71, 134]. Briefly, sequential illumination was provided at 470 nm, 530 nm, and 625 nm; reflected light in each channel was collected by a lens (focal length = 75mm, NMV-75M1, Navitar), split by a 580 nm dichroic (FF580-FDi02-t3-25×36, Semrock) into two channels, one filtered by 500 nm long pass (FF01-500/LP-25, Semrock – blocks FAF excitation, passes FAF emission and 530 nm Hb reflectance), the other by a 593 nm long pass (FF01-593/LP-25, Semrock - blocks jRGECO1a excitation, passes jRGECO1a emission and 625 nm Hb reflectance). The two channels were detected separately via two CMOS cameras (Zyla 5.5, Andor). Data were cropped to 1024×1024 pixels, and binned to 512×512 to achieve a frame rate of 100Hz in each camera, with all contrasts imaged at 25 Hz. The LEDs and camera exposures were synchronized and triggered via a data acquisition card (PCI-6733, National Instruments) using MATLAB R2019a (MathWorks).

Mice were head-fixed under the imaging system objective using an aluminum bracket attached to a skull-mounted Plexiglas window. Prior to data acquisition in the awake state, mice were acclimated to head-fixation over several days.

The mouse’s body was supported by a felt pouch suspended by optical posts (Thorlabs). Resting state imaging was performed for 10 minutes in each mouse. Before each imaging run, dark counts were imaged for each mouse for 1 second with all LEDs off in order to remove background sensor noise.

##### Preprocessing

Images were spatially normalized, downsampled to 128×128 pixels, co-registered, and affine-transformed to the Paxinos atlas, temporally detrended (5^th^ order polynomial fit) and spatially smoothed (5×5 pixel Gaussian filter with standard deviation of 1.3 pixels) [135]. Changes in 530 nm, and 625 nm reflectance were interpreted using the modified Beer-Lambert law to calculate changes in oxy- and deoxy-hemoglobin concentration [71].

Image sequences of fluorescence emission detected by CMOS1 (i.e., uncorrected FAF) and CMOS2 (i.e., uncorrected jRGECO1a) were converted to percent change (dF/F) by dividing each pixel’s time trace by its average fluorescence over each imaging run. Absorption of excitation and emission light for each fluorophore due to hemoglobin was corrected as outlined in [136]. From the face videography we derived scalar indices of pupil size via DeepLabCut software [137] and whisker motion via the Lucas-Kanade optical flow method [138] applied to and subsequently averaged across five manually selected data points on the whiskers. The resulting pupil diameter, whisker motion, and widefield time series were bandpass filtered between 0.01 *< f <* 0.2 Hz in order to distinguish the hypothesized spatiotemporal process from distinct phenomena occurring at higher frequencies (e.g., slow waves) [53], and to accommodate finite scan duration 10 minutes. For visualization only—namely, to view the Allen atlas boundaries in register with cortical maps—we manually aligned the Paxinos-registered data from one mouse to a 2D projection of the Allen Mouse Common Coordinate Framework v3 [139] using four anatomical landmarks: the left, right, and midline points at which the anterior cortex meets the olfactory bulbs, and an additional midline point at the base of retrosplenial cortex [9, 140]. Coordinates obtained from this transformation were used to overlay the CCF boundaries onto the Paxinos-registered cortical maps for all mice.

#### Dataset 2: Allen Brain Observatory

We additionally analyzed recordings obtained from awake mice and publicly released via the Allen Brain Observatory Neuropixels Visual Coding project [104]. These recordings include eye-tracking, running speed (estimated from running wheel velocity), and high-dimensional electrophysiological recordings obtained with Neuropixels probes [105]. We restricted analyses to the “Functional Connectivity” stimulus set, which included a 30-minute “spontaneous” session with no overt experimental stimulation. Data were accessed via the Allen Software Development Kit (SDK) (https://allensdk.readthedocs.io/en/latest/). From 26 available recording sessions, we excluded four with missing or compromised eye-tracking data, one lacking local field potential (LFP) data, and five with locomotion data indicating a lack of immobile periods or appearing otherwise anomalous, thus leaving a total of 16 recordings for downstream analyses.

##### Preprocessing

From the electrophysiological data we derived estimates of population firing rates and several quantities based upon the LFP. We accessed spiking activity as already extracted by Kilosort2 [7]. Spikes were binned in 1.2 s bins as previously [7], and a mean firing rate was computed across all available units from neocortical regions surpassing the default quality control criteria. Neocortical LFPs were used to compute band-limited power (BLP) within three canonical frequency bands: gamma (40 – 100 Hz), “alpha” (3 − 6 Hz) and delta (0.5 − 4 Hz). Specifically, we downsampled and low-passed the signals with a 100 Hz cutoff, computed the spectrogram (*scipy.signal.spectrogram* [141]) at each channel in sliding windows of 0.5 s with 80% overlap, and averaged spectrograms across all channels falling within “VIS” regions (including primary and secondary visual cortical areas). BLP was then computed by integrating the spectrogram within the frequency bands of interest and normalizing by the total power. A similar procedure was applied to LFP recordings from the hippocampal CA1 region to derive estimates of hippocampal theta (5 − 8 Hz) and delta (0.5 − 4 Hz) BLP. Hippocampal sharp-wave ripples were detected on the basis of the hippocampal CA1 LFP via an automated algorithm [142] following previously described procedures [143]. In general, extreme outlying values (generally artifact) were largely mitigated through median filters applied to all observables, whereas an additional thresholding procedure was used to interpolate over large negative transients in running speed. Finally, all time series were downsampled to a sampling rate of 20 Hz and filtered between 0.01 *< f <* 0.2 Hz to facilitate integration with Dataset 1.

### Data analysis

#### State-space framework

Formally, we consider the generic state-space model:

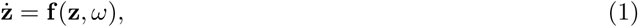

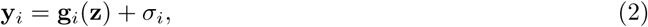

where the nonlinear function **f** determines the (nonautonomous) flow of the latent (i.e., unobserved), vector-valued arousal state **z** = [*z*_1_*, z*_2_*,…, z_m_*]^T^ along a low-dimensional attractor manifold M, while *ω*(*t*) reflects random (external or internal) perturbations decoupled from **f** that nonetheless influence the evolution of **z** (i.e., “dynamical noise”). We consider our *N* observables 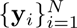 as measurements of the arousal dynamics, each resulting from an observation equation **g***_i_*, along with other contributions *σ_i_*(*t*) that are decoupled from the dynamics of **z** (and in general are unique to each observable). Thus, in this framing, samples of the observable **y***_i_* at consecutive time points *t* and *t* + 1 are linked only through the evolution of the latent variable **z** as determined by **f**. Note that **f** is presumed to embody various causal influences spanning both brain and body (and the feedback between them), whereas *measurements* (of either brain or body) **y***_i_* do not contribute to these dynamics. This framework provides a formal, data-driven approach to parsimoniously capture the diverse manifestations of arousal dynamics, represented by the proportion of variance in each observable **y***_i_* that can be modeled purely as a time-invariant mapping from the state-space of **z**.

### State-space reconstruction: time delay embedding

Our principal task is to learn a mapping from arousal-related observables to the multidimensional space where the hypothesized arousal process evolves in time. To do this, we take advantage of Takens’ embedding theorem from dynamical systems theory, which has been widely used for the purpose of nonlinear state-space reconstruction from an observable. Given *p* snapshots of the scalar observable *y* in time, we begin by constructing the following Hankel matrix **H** ∈ R*^d^*^×(^*^p^*^−^*^d^*^+1)^:

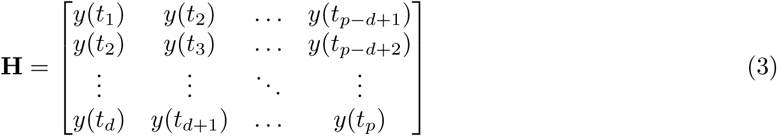

for *p* time points and *d* time delays. Each column of **H** represents a short trajectory of the scalar observable *y*(*t*) over *d* time points, which we refer to as the *delay vector* **h**(*t*). These delay vectors represent the evolution of the observable within an augmented, *d*−dimensional state-space.

We initially construct **H** as a high-dimensional (and rank-deficient) matrix to ensure it covers a sufficiently large span to embed the manifold. We subsequently reduce the dimensionality of this matrix to improve conditioning and reduce noise. Dimensionality reduction is carried out in two steps—first, through projection onto an orthogonal set of basis vectors (below), and subsequently through nonlinear dimensionality reduction via a neural network (detailed in the next section).

For the initial projection, we note that the leading left eigenvectors of **H** converge to Legendre polynomials in the limit of short delay windows [144]. Accordingly, we use the first *r* = 10 discrete Legendre polynomials as the basis vectors of an orthogonal projection matrix:

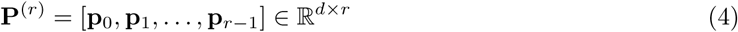

(polynomials obtained from the *special.legendre* function in SciPy [141]). We apply this projection to the Hankel matrix constructed from a pupil diameter timecourse. The resulting matrix 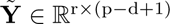 is given by:

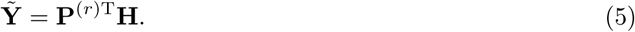

Each column of this matrix, denoted 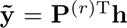, represents the projection of a delay vector **h**(*t*) onto the leading *r* Legendre polynomials. The components of this state vector, 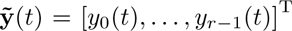, are the individual *r Legendre coordinates* [144], which form the input to the neural network encoder.

The dimensionality of **H** is commonly reduced through its singular value decomposition (SVD) (e.g., [70, 145]). In practice, we find that projection onto Legendre polynomials yields marginal improvements over SVD of **H**, particularly in the low-data limit [144]. More importantly, the Legendre polynomials additionally provide a universal, analytic basis in which to represent the dynamics; this property is exploited for comparisons across mice and datasets.

Choice of delay embedding parameters was guided on the basis of autocorrelation time and attractor reconstruction quality (i.e., unfolding the attractor while maximally preserving geometry), following decomposition of **H**. The number of delay coordinates was guided based upon asymptoting reconstruction performance using a linear regression model. For all main text analyses, the Hankel matrix was constructed with *d* = 100 time delays, with each row separated by Δ*t* = 3 time steps (equivalently, adopting **H** as defined in Eq. 3, we set *d* = 300 and subsampled every third row). As all time series were analyzed at a sampling frequency of 20 Hz, this amounts to a maximum delay window of 100 × 3 × .05 = 15 seconds.

For all modeling, pupil diameter was first shifted in time to accommodate any physiological delay between the widefield signals and the pupil. For each widefield modality, this time shift was selected as the abscissa corresponding to the peak of the cross-correlation function between the pupil and the mean widefield signal. For the leave-one-out pipeline, we used the median time shift across the six mice in the training set.

### State-space mappings: variational autoencoder

To interrelate observables through this state-space framework (and thus, through arousal dynamics), we ultimately seek to approximate the target functions

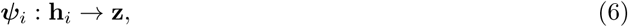

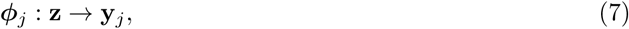

where **h***_i_*(*t*) represents the delay vectors (columns of the Hankel matrix constructed from a scalar observable *y_i_*), **z**(*t*) ∈ R*^m^* is the low-dimensional representation of the latent arousal state, and **y***_j_*(*t*) is a second observable. A variety of linear and nonlinear methods can be used to approximate these functions (e.g., see [86]). Our primary results are obtained through a probabilistic modeling architecture based upon the variational autoencoder (VAE) [91, 146].

Briefly, a VAE is a deep generative model comprising a pair of neural networks (i.e., an encoder and decoder) that are jointly trained to map data observations to the mean and variance of a latent distribution (via the encoder), and to map random samples from this latent distribution back to the observation space (decoder). Thus, the VAE assumes that data observations *y* are taken from a distribution over some latent variable *z*, such that each data point is treated as a sample 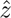 from the prior distribution *p_θ_*(*z*), typically initialized according to the standard diagonal Gaussian prior, i.e., *p_θ_*(*z*) = N (*z*|0, **I**). A variational distribution *q_φ_*(*z*|*y*) with trainable weights *φ* is introduced as an approximation to the true (but intractable) posterior distribution *p*(*z*|*y*).

As an autoencoder, *q_φ_*(*z*|*y*) and *p_θ_*(*y*|*z*) encode observations into a stochastic latent space, and decode from this latent space back to the original observation space. The training goal is to maximize the marginal likelihood of the observables given the latent states *z*. This problem is made tractable by instead maximizing an evidence lower-bound (ELBO), such that the loss is defined in terms of the negative ELBO:

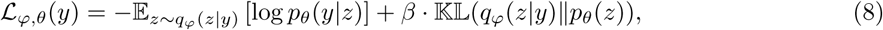

where the first term corresponds to the log-likelihood of the data (reconstruction error), and the Kullback-Leibler (KL) divergence term regularizes the distribution of the latent states *q_φ_*(*z*|*y*) to be close to that of the prior *p_θ_*(*z*). Under Gaussian assumptions, the first term is simply obtained as the mean squared error, i.e., 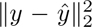. The KL-term is weighted according to an additional hyperparameter *β*, which controls the balance between these two losses [147, 148]. Model parameters (*φ* and *θ*) can be jointly optimized via stochastic gradient descent through the reparameterization trick [146].

In the present framework, rather than mapping back to the original observation space, we use the decoder *p_θ_*(*y*|*z*) to map from **z** to a new observation space. In this way, we use a probabilistic encoder and decoder, respectively, to approximate our target functions ***ψ***(**h**) (which is partly constituted by the intermediate projection **P**^(^*^r^*^)^, via Eq. 5) and ***ϕ***(**z**). Notably, VAE training incorporates stochastic perturbations to the latent representation, thus promoting discovery of a smooth and continuous manifold. Such a representation is desirable in the present framework, such that changes within the latent space (which is based on the delay coordinates) are smoothly mapped to changes within the observation spaces [149]. After training, to generate predicted trajectories, we simply fed the mean output from the encoder (i.e., *q*(*µ_z_*|*y*)) through the decoder to (deterministically) generate the maximum a posteriori (MAP) estimate.

### Model architecture and hyperparameters

VAE models were trained with the Adam optimizer [150]; see Table 1 for details specific to each dataset. Across datasets, the encoder comprised a single-layer feedforward neural network with tanh activations, which transformed the vector of leading Legendre coordinates 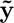 (Eq. (5)) to (the mean and log variance of) an *m*-dimensional latent space (*m* = 4 unless otherwise stated). The latent space Gaussian was initialized with a noise scale of *σ* = .1 to stabilize training. The decoder from this latent space included a single-layer feedforward neural network consisting of 10 hidden units with tanh activations, followed by a linear readout layer matching the dimensionality of the target signal. During training, the KL divergence weight was gradually annealed to a factor of *β* over the first ∼ 15% of the training epochs to mitigate posterior collapse [147, 151]. Hyperparameters pertaining to delay embedding and the VAE were primarily selected by examining expressivity and robustness within the training set of a subset of mice. Within each modeling pipeline/dataset, hyperparameters were kept fixed across all mice and modalities to mitigate overfitting.

**Table 1:** Neural network hyperparameters across modeling pipelines and datasets.

| Hyperparameter | Leave-one-out | Within-mouse | Allen mouse | Lorenz |
| --- | --- | --- | --- | --- |
| Latent dimensions ($z_n$) | 4 | 4 | 2 | 6 |
| Learning rate (LR) | 0.001 | 0.01 | 0.001 | 0.01 |
| Epochs | 200 | 200 | 30 | 200 |
| KL weight ($\beta$) | 0.1 | 1 | 0.1 | 0.1 |
| KL anneal steps | 2000 | 200 | 2000 | 2000 |
| Batch size | 1000 | 1000 | 1000 | 1000 |

### Model training pipeline

For all main analyses, we implemented an iterative leave-one-out strategy in which candidate models were trained using 10-minute recordings concatenated across six of the seven mice, and subsequently tested on the held-out 10-minute recording obtained from the seventh mouse. To make model training efficient, we employed a randomized singular value decomposition (SVD) approach [68, 152]. Thus, for each iteration, we began by pre-standardizing (to zero mean and unit variance) all datasets (“standardization 1”) and aligning in time following delay embedding of the pupil measurements (thus removing the initial time points of the widefield data for which no delay vector of pupil values is available). Next, we lag-adjusted the (delay embedded) pupil and widefield time series from each mouse according to the median lag (across the six training mice) derived from each training mouse’s cross-correlation function between their pupil time series and their mean widefield signal. Next, we concatenated the six training datasets and computed the SVD of the concatenated widefield time series 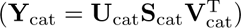 to obtain group-level spatial components **V**_cat_. Data were projected onto the leading 10 components (i.e., the first 10 columns of **V**_cat_) and standardized once more (“standardization 2”), and training proceeded within this reduced subspace. Note that this group-level subspace excludes data from the test mouse, such that **V**_cat_ differs for each iteration of the leave-one-out pipeline.

For the held-out mouse, the pre-standardized pupil diameter (i.e., having undergone “standardization 1” above) was standardized once more using the training set parameters (“standardization 2”), and used to generate a prediction within the group-level 10-dimensional subspace. This prediction was then inverse-transformed and unprojected back into the high-dimensional image space using the training set’s standardization and projection parameters. All model evaluations were performed following one final inverse transformation to undo the initial pre-standardization (standardization 1), enabling evaluation in the original dF/F units. The pre-standardization parameters were not predicted from the training data; including the pre-standardization as part of the model would make model performance sensitive to the robustness of the relationship between dF/F values and raw pupil units, which is not scientifically relevant in this context. Finally, any subsequent projections used to evaluate model performance in terms of PCs (as in Fig. 2B) utilized mouse-specific SVD modes (denoted simply as **V**), rather than the group modes (**V**_cat_) used during training.

In addition to the leave-one-out pipeline, we implemented a separate, within-subject pipeline in which we trained models on the first 5 minutes of data for each mouse and report model performance on the final 3.5 minutes of the 10-minute session. This training pipeline involved prediction of the full-dimensional images, without the intermediate SVD used in the leave-one-out pipeline. All results pertaining to this modeling pipeline are contained in Fig. S9.

### Model evaluation

For analyses reporting “Total variance” explained (Figs 2A and 3C, upper), we report the *R*^2^ value over the full 128 × 128 image, computed across all held-out time points. *R*^2^ was computed as the coefficient of determination, evaluated over all pixels *n* and temporal samples *t* (i.e., the total variance):

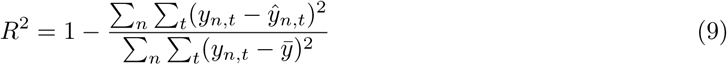

where *y_n,t_* and *ŷ_n,t_* are, respectively, the true and predicted values of **y** at the *n*-th pixel and *t*-th time point, and 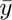 is the global mean. In practice, this value was computed as the pixel-wise weighted average of *R*^2^ scores, with weights determined by pixel variance (computed using the built-in function *sklearn.metrics.r2 score*, with “multioutput” set to “variance-weighted”).

To examine variance explained beyond the first principal component (Figs. 2B and 3C, lower), we computed the (randomized) SVD of the widefield data for each mouse (**Y** = **USV**^T^) and, after training, projected the original and reconstructed widefield data onto the top *N* = 200 spatial components excluding the first (i.e, **YV**_2:_*_N_*) prior to computing *R*^2^ values. The matrices from this SVD were used for all analyses involving projection onto the leading PCs.

For the shuffled control analysis (Fig. S16), we examined total variance explained in widefield data from the test set (as in Fig. 2A), except that for each mouse we swapped the original pupil diameter timecourse with one from each of the other six mice (i.e., we shuffled pupil diameter and widefield calcium data series across mice).

### Dynamic mode decomposition

Dynamic mode decomposition (DMD) [153–155] is a data-driven method to dimensionally reduce time series measurements into a superposition of spatiotemporal modes. Given the time series matrix **Y** = {**y**(*t*_1_), **y**(*t*_2_)*,…,* **y**(*t_p_*)}, where **y**(*t_i_*) represents the system state vector at the *i*th time point, the data are split into two matrices, **Y**^(1)^ = {**y**(*t*_1_)*,…,* **y**(*t_p_*_−1_)} and **Y**^(2)^ = {**y**(*t*_2_)*,…,* **y**(*t_p_*)}. DMD seeks an approximate linear mapping **A** such that

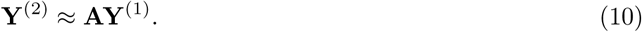

Theoretically, the operator **A** represents a low-rank approximation to an infinite-dimensional linear operator—namely, the Koopman operator—associated with nonlinear systems [69, 155], motivating use of DMD even in the context of nonlinear dynamics. This low-rank approximation is obtained via SVD of **Y**^(1)^ and **Y**^(2)^. The measurement data can thus be approximated in terms of the spectral decomposition of **A**:

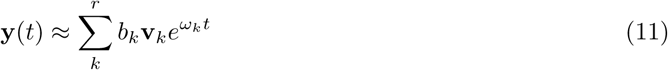

where the eigenvectors **v***_k_* give the dominant spatial modes, the eigenvalues *ω_k_* give the dominant temporal frequencies, and the coefficients *b_k_* determine the amplitudes.

For each mouse, we applied DMD separately to three data matrices: the original widefield calcium images, and the predictions generated through both the “No embedding” model and the “Latent model”. Here again, for numerical efficiency, we incorporated randomized SVD as part of the randomized DMD algorithm [156] implemented in *pyDMD* [157, 158].

Following application of DMD, we constructed spatial phase maps from the leading eigenvector of each **A** matrix. Specifically, given the leading eigenvector **v**_1_ = [*v*_1_*, v*_2_*,…, v_n_*]^T^ ∈ C*^n^*, we computed the phase at each pixel *i* as *θ_i_* = arg(*v_i_*), where arg(·) denotes the complex argument of each entry, and *v_i_* represents the value of the eigenvector **v**_1_ at the *i*-th pixel. Note that the “No embedding” model predictions yielded exclusively real-valued eigenvectors, as these model predictions were fundamentally rank-one (i.e., they reflect linear transformations of the scalar pupil diameter). Accordingly, as the complex argument maps negative real numbers to 0 and positive real numbers to *π*, phase maps obtained from the “No embedding” model predictions were restricted to these two values.

We used the circular correlation coefficient to quantify the similarity between the phase map obtained from the original dataset and the phase map obtained from either of the two model predictions. Specifically, let *θ_i_* and 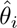 represent the phase at spatial location *i* in the phase maps of the original and reconstructed datasets. The circular correlation *ρ*_circ_ between the two phase maps was computed as [159]:

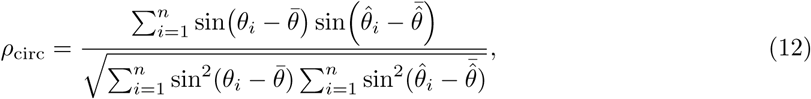

where 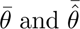 are the mean phase angles (i.e., the mean directions) of the respective maps. The circular correlation is bounded between −1 and 1, where 1 indicates perfect spatial phase alignment between the datasets, and −1 indicates an inverted pattern. However, because the eigenvectors are only determined up to a complex sign, Fig. 2C reports the absolute value this quantity, i.e., |*ρ*_circ_|.

### Clustering and decoding analysis

We used Gaussian mixture models (GMMs) to assign each (variance-normalized) image frame to one of *k* clusters (parameterized by the mean and covariance of a corresponding multivariate Gaussian) in an unsupervised fashion. This procedure enables assessment of the spatial specificity of cortical patterns predicted by arousal dynamics, with increasing number of clusters reflecting increased spatial specificity.

Briefly, a GMM models the observed distribution of feature values *x* as coming from some combination of *k* Gaussian distributions:

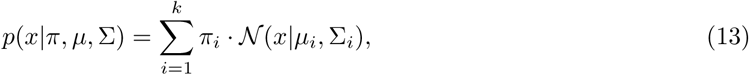

where *µ_i_* and Σ*_i_* are the mean and covariance matrix of the *i*th Gaussian, respectively, and *π_i_* is the probability of *x* belonging to the *i*th Gaussian (i.e., the Gaussian weight). We fit the mean, covariance, and weight parameters through the standard expectation-maximization algorithm (EM) as implemented in the GMM package available in *scikit-learn* [160]. This procedure results in a posterior probability for each data point’s membership to each of the *k* Gaussians (also referred to as the “responsibility” of Gaussian *i* for the data point); (hard) clustering can then be performed by simply assigning each data point to the Gaussian cluster with maximal responsibility.

Images were spatially normalized such that the (brain-masked) image at each time point was set to unit variance (thus yielding a “spatial sign” [161]). Then, for each mouse and each number of clusters *k*, a GMM with full covariance prior was fit to the normalized image frames to obtain a set of “ground truth” cluster assignments. After training, this GMM was subsequently conditioned on the reconstructed rather than original image frames to obtain maximum a posteriori cluster assignments associated with the “No embedding” or “Latent model” predictions. These cluster assignments were compared to the ground truth assignments, with accuracy computed as the proportion of correctly identified cluster labels.

The preceding algorithm effectively clusters the dimensionally-reduced image frames on the basis of their cosine distance (technically, their *L*_2_-normalized Euclidean distance). Because the first PC tends to explain the vast majority of the variance, this Euclidean-based distance is largely determined by distance along the leading dimension. This analysis choice suffices for the primary purpose of the analysis— namely, to assess correspondence between the original and reconstructed data at the level of individual time points. However, in the context of multivariate data, it is common to also normalize the data *features* prior to clustering, thus equally weighting each (spatial) dimension of the data. To examine this clustering alternative, we performed *feature-wise* normalization of each spatial dimension following projection of the (*sample-wise*) normalized image frames onto the top three PCs. This essentially constitutes a whitening transformation of the image frames, with Euclidean distance in this transformed space effectively approximating the Mahalanobis distance between image frames. Clustering results from this procedure are shown in Fig. S11. For these supplementary analyses, which we also extend to higher cluster numbers, we employ a spherical (rather than full) covariance prior as additional regularization.

### Hidden Markov model

We additionally used hidden Markov models (HMMs) to capture the temporal dynamics within the original and reconstructed widefield calcium measurements. Briefly, under a Gaussian HMM, the observations {**y**(*t*_1_), **y**(*t*_2_)*,…,* **y**(*t_p_*)} are assumed to be generated by sequence of latent discrete states *z_t_* ∈ {1*,…, K*}, with the full generative model including:

1. A transition matrix **A** ∈ R*^K^*^×^*^K^*, where *A_ij_* = *P* (*z_t_*_+1_ = *j*|*z_t_* = *i*) represents the probability of transitioning from state *i* to state *j*.
2. An initial state distribution, represented as a vector of state probabilities ***π*** ∈ R*^K^* (with *π_i_* = *P* (*z*_1_ = *i*)).
3. Emission distributions, such that each observation **y***_t_* is modeled as a multivariate Gaussian conditioned on the latent state *z_t_*—i.e., **y***_t_* ∼ N (***µ****_z_,* **Σ***_z_*), where ***µ****_z_* and **Σ***_z_* denote the state-specific mean and covariance, respectively.

The model is thus parameterized by 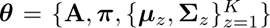. We estimated these parameters via an EM algorithm (Baum-Welch), as implemented in *hmmlearn* [162].

Prior to HMM fitting, the widefield images were projected onto the three leading PCs and normalized along each dimension (similar to preprocessing for the GMM with whitening above, but without the initial normalization of each individual image frame). HMMs were fit to these data for a range of state numbers (*K* = 2, 3, 4, 5, 6).

To ensure a robust fit, we initialized the state probabilities to be uniform, 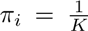, and initialized the transition matrix with probabilities of .8 along the diagonal (i.e., within-state transitions or “stay” probabilities) and .2*/K* for all off-diagonal elements (i.e., between-state transitions).

An HMM was first fit to the original data (yielding a “ground truth” model), and then separate HMMs were fit to the reconstructions. In the latter cases, the emission parameters 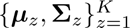 were fixed to the values obtained from the ground truth model, so that only the transition matrix **A** and the initial state probabilities ***π*** were updated.

For each mouse and each choice of *K* states, we computed three evaluation metrics:

1. The log-likelihood of the observed data under the fitted HMMs.
2. Decoding accuracy, computed as the agreement between the latent states inferred from the ground truth HMM and those predicted by the HMMs fit to the reconstructed datasets.
3. Transition matrix similarity, which we computed as the row-wise KL-divergence between the ground truth and reconstructed transition matrices:

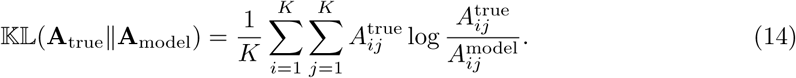

For numerical stability in computing KL divergence, transition probabilities below a threshold of *ɛ* = 10^−6^ were set to zero and each row was renormalized to maintain stochasticity, i.e.,

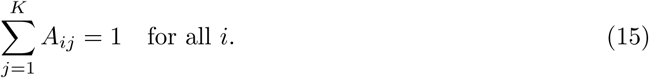

### Multi-dataset integration

To extend our framework to an independent set of observables, we trained our architecture on concatenated time series from the Allen Institute Brain Observatory dataset (*N* = 16 mice). For visualization purposes, we sought to learn a group-level two-dimensional embedding (i.e., **z** ∈ R^2^) based upon delay embedded pupil measurements. After group-level training, we froze the weights of the encoder and retrained the decoders independently for each mouse, enabling individual-specific predictions for the Allen Institute mice from a common (i.e., group-level) latent space. We additionally applied these encoder weights to delay embedded pupil observations from the original “WashU” dataset, once again retraining mouse-specific decoders to reconstruct observables from the common latent space.

Once trained, this procedure results in the generative model *p*(**y***_i_,* **z**), which we use to express the posterior probability of each observable **y***_i_* as a function of position within a common 2D latent space **z** (note that this formulation does not seek a complete generative model capturing all joint probabilities among the observables). This allows us to systematically evaluate the conditional expectation (E(**y***_i_*|**z**)) for each observable over a grid of points in the latent space, which was regularized to be continuous (roughly, an isotropic Gaussian [91]). Fig. 4 represents this expectation, averaged over decoder models trained for each mouse.

## Acknowledgements

We wish to thank the Allen Institute for the publicly available data used in this study. We also thank Trevor Voss for assistance with the optical setup. The mouse cartoon in Figs. 1 & 4 is adapted from [163], obtained from SciDraw.io.

## Data availability statement

The multimodal widefield datasets collected for this study will be deposited in a public repository upon publication. Data from the Allen Brain Observatory can be accessed via the Allen SDK (https://allensdk.readthedocs.io/en/latest/).

## Code availability statement

All analysis code for this study is publicly available at https://github.com/ryraut/arousal_dynamics.

## Supplementary Text

### Toy example: Stochastic Lorenz system

We demonstrate the efficacy of our procedure on the three-dimensional Lorenz system, a canonical low-dimensional nonlinear dynamical system. Specifically, we simulated the following stochastic system:

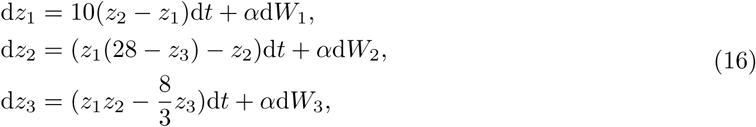

where coefficients reflect standard parameter choices, and *α* is the scaling parameter applied to three independent Wiener processes (*W*_1_*, W*_2_*, W*_3_) acting on the three state variables.

For all simulations, the observation functions *g_n_* are simply identity functions, the scalar observable *y*_1_ (by analogy, pupil diameter) is a function of the state variable *z*_2_, and the multidimensional observable **y**_2_ (by analogy, a brain image timecourse) is a function of the other two state variables, [*z*_1_*, z*_3_]^T^). We considered multiple levels of dynamical noise *ω*(*t*), which was modeled as a multidimensional Wiener process scaled by the parameter *α*, and multiple levels of the observation noises *σ*_1_ and *σ*_2_, which were modeled as unit Gaussians multiplied by a scaling factor *β*. Thus, the stochastic Lorenz governing equations (Eq. 16) were coupled to the observation functions:

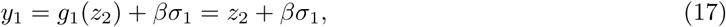

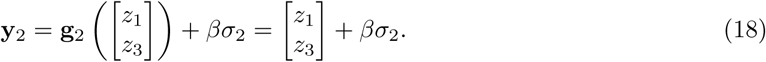

We sought to reconstruct the dynamics of the 2D observable **y**_2_ on the basis of a time delay embedding of the scalar observable *y*_1_. As in the main text analyses, we assessed reconstruction accuracy in test data as a function of whether delay coordinates and/or nonlinearities were included in the model.

Lorenz trajectories were simulated with Δ*t* = .01 and *N* = 10000 time steps. The first 6000 time steps were included in the training set, and the model was tested on the final 3000 time steps. Stochastic trajectories (i.e., where *α* ≠ 0) were simulated with the standard Euler-Maruyama integration scheme.

Fig. S3 demonstrates the efficacy of this procedure across a range of values for the dynamical noise and observation noise (controlled by parameters *α* and *β*, respectively). With no dynamical noise or observation noise, the “Latent model” enabled nearly perfect prediction of the multidimensional observable on the basis of the scalar observable *y*_1_ and its dynamics. Delay embedding and nonlinearities were generally both crucial to optimal performance. Across all models, performance gradually degraded with increasing levels of dynamical noise (Fig. S3B-C). However, notably, performance could remain high even in the presence of dynamical noise that introduces obvious distortion to the latent trajectories. This is because the noise is broadcast throughout the system, and is thus manifested in the dynamics of both observables.

On the other hand, with increasing levels of observation noise (increasing *β*), which is derived from Gaussian fluctuations unique to *y*_1_ and **y**_2_, the proportion of predictable variance in **y**_2_ rapidly decreased (Fig. S3 D-E). This is because variance in **y**_2_ became increasingly dominated by fluctuations that are not manifested in *y*_1_, precluding the ability to model **y**_2_ via a time-invariant mapping from a state-space reconstruction based on *y*_1_.

In the context of arousal dynamics, observation noise in our framework includes genuine measurement noise (e.g., imaging artifacts) as well as physiological sources of variance that are conditionally independent of arousal (e.g., fluctuations resulting from sensory-evoked neural activity, or processes other than “arousal” that are hypothetically reflected in pupil diameter [130]). Accordingly, if model performance is poor, this could reflect either genuine measurement noise or the predominance of physiological processes that are not common to both observables. By contrast, the strong performance of our method is an indication of the relatively minor contribution from both of these sources of observation noise in comparison to a dominant, shared latent process.

### Universal embeddings via Legendre polynomials

The leading left eigenvectors of **H** approximate eigenfunctions of the Koopman operator K, which advances measurements of a dynamical system [70]. Importantly, the eigenfunctions of K define a subspace invariant to the dynamics. These eigenfunctions can be used to parameterize the manifold based on linearization around a fixed point, extended throughout the basin of attraction [164]. Thus, whereas Results in Fig. 4 reflect mappings from a 2D latent space obtained through a neural network, these properties of Koopman eigenfunctions imply the possibility of using the leading Legendre coordinates (Eq. (5)) as intrinsic, analytical coordinates for parameterizing the arousal manifold M.

## Supplementary Figures

**Figure S1:**
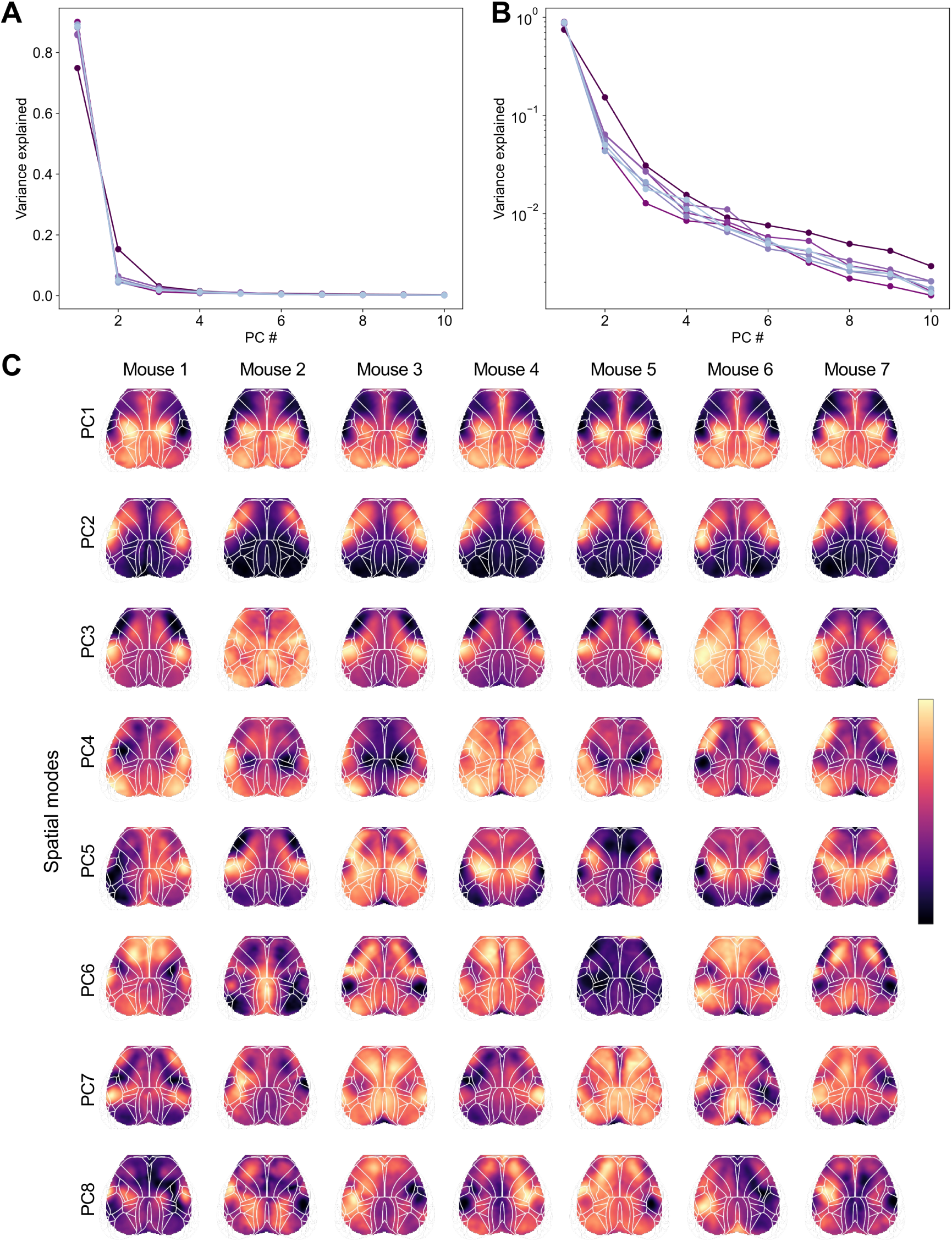
SVD results for widefield calcium images. **A-B** Scree plots for each of the seven mice, displayed on a linear (A) or log-transformed (B) y-axis. For all mice, images were dominated by the first principal component. **C** Spatial topographies corresponding to the top 8 PCs (arbitrary units). As the sign of each PC is arbitrary, some PCs were sign-flipped to improve visual correspondence across mice.

**Figure S2:**
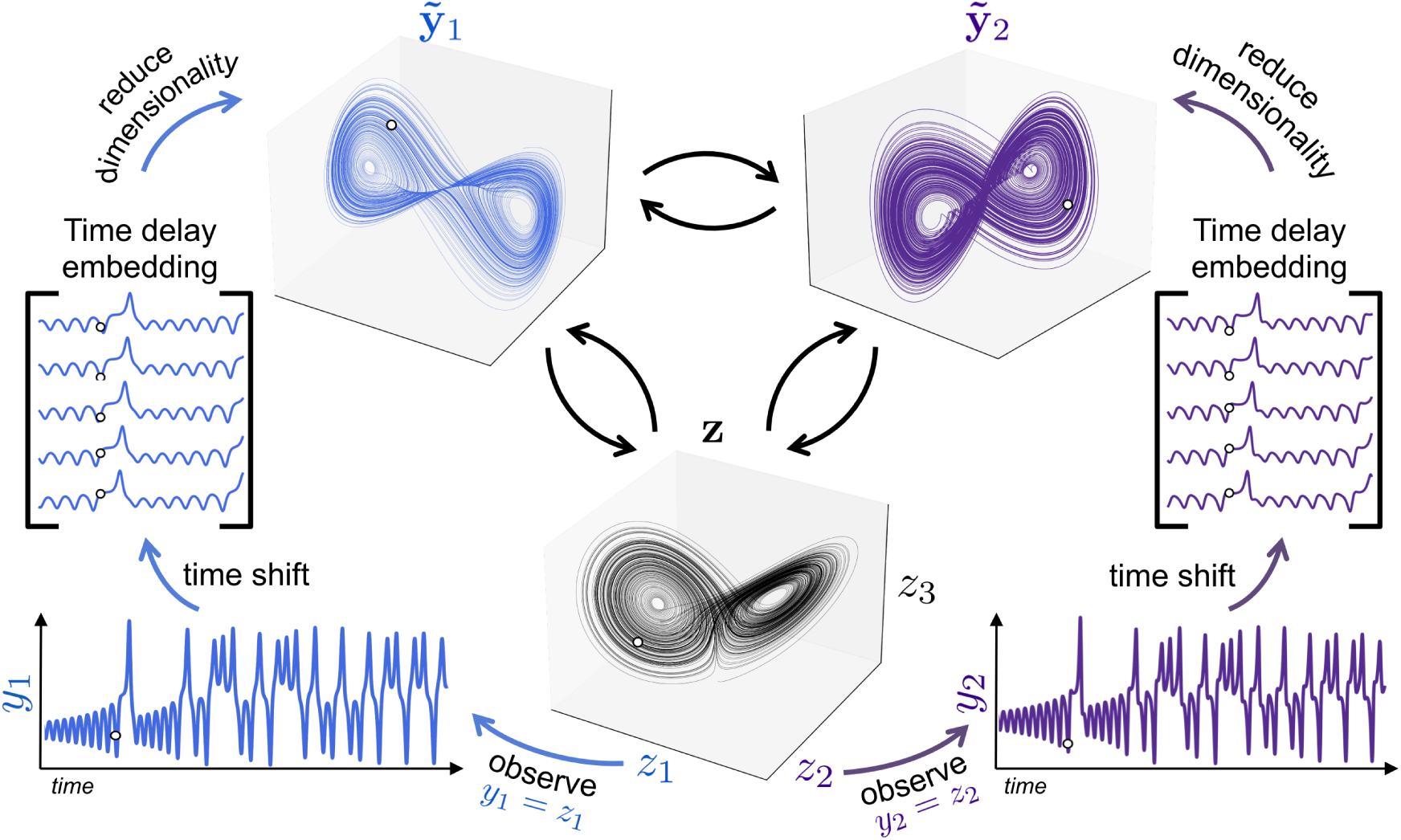
Illustration of shadow manifolds in the context of the 3D Lorenz system. The white dot in each plot represents the same point in time (i.e., the same dynamical state) from several different views (or embeddings). Here, the measurements *y*1 and *y*2 are simply taken to be the state variables *z*1 and *z*2 from the original system **z**(*t*). Each of these observations can be stacked with time-shifted copies of itself to obtain a (high-dimensional) time delay embedding that is diffeomorphic to the original attractor. These embeddings can be dimensionally reduced to obtain low-dimensional “shadow manifolds” that (approximately) preserve the topology of the original system. Pairs of black arrows indicate relation via a diffeomorphic mapping. These relations clarify the motivation for modeling one observable from a dynamical system (e.g., *y*2) as a function of a time delay embedding of a second observable (e.g., the augmented observable 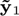). See Fig. ^S3^ for demonstration of our computational framework applied to the Lorenz system.

**Figure S3:**
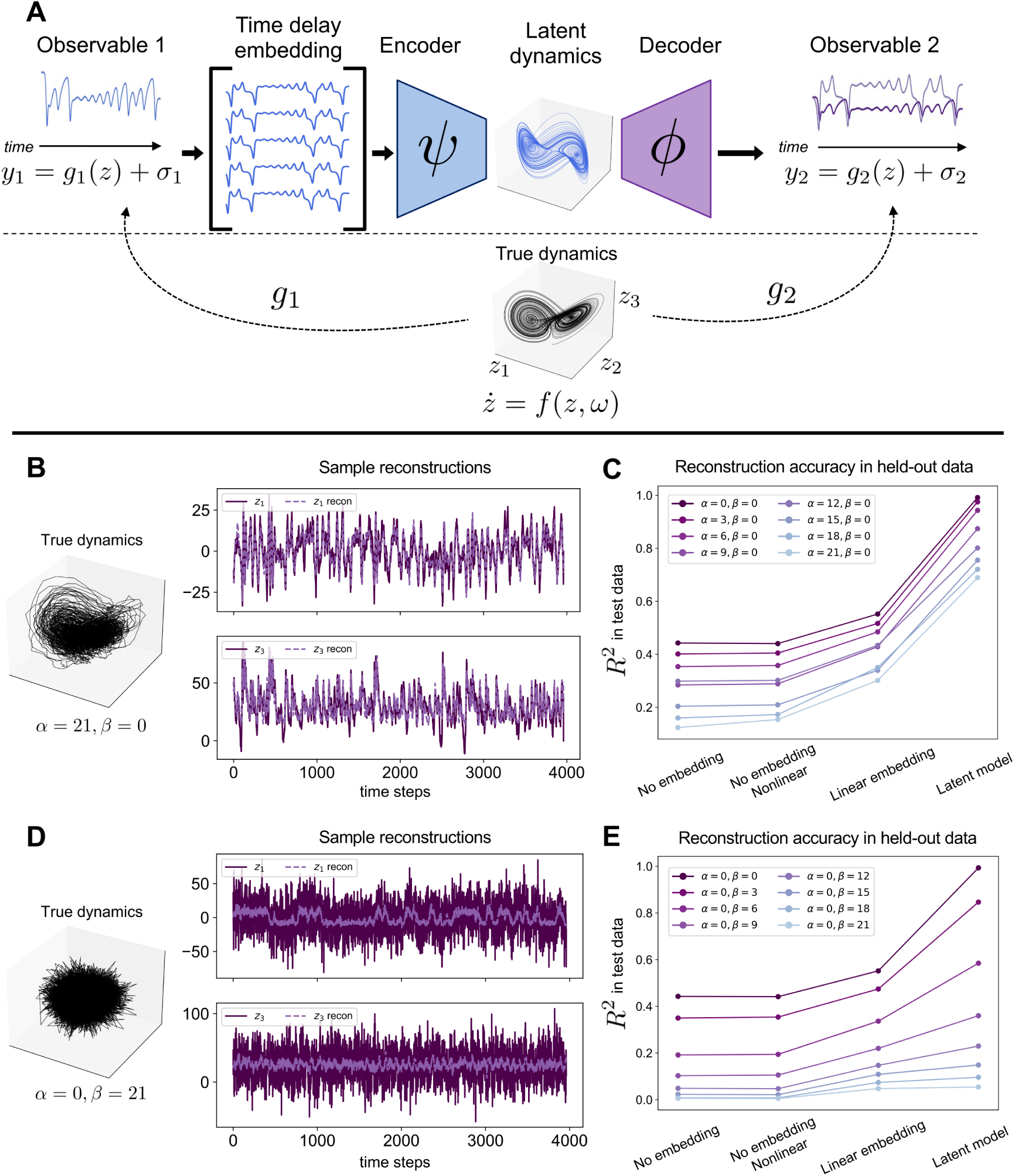
Toy demonstration of delay embedding for multimodal cross-prediction in the presence of dynamical and observation noise. **A** Delay embedding framework applied to a stochastic version of the 3D Lorenz system (compare to Fig. 1C). Here, the observation functions *g*1 and *g*2 are defined as identity functions. Across simulations, *y*1 is taken to be a function of *z*2, whereas **y**2 is a 2D function of *z*1 and *z*3 . We aim to model **y**2 according to *y*1 (that is, a noisy measurement of a single state variable, *z*2), across different levels of dynamical noise and observation noise. *f* corresponds to the stochastic Lorenz system defined in Eq. 16. **B-C** Reconstruction accuracy across multiple levels of dynamical noise, scaled by the parameter *α*. B illustrates sample reconstructions of the state variables *z*1 and *z*3 from the observable *y*1 at the highest dynamical noise level tested (*α* = 21). **D-E** Same as B-C, but across multiple levels of observation noise, scaled by the parameter *β*. D illustrates sample reconstructions for the state variable time series at the highest noise level of observation noise tested (*β* = 21). See Supplementary Text for further details and interpretation.

**Figure S4:**
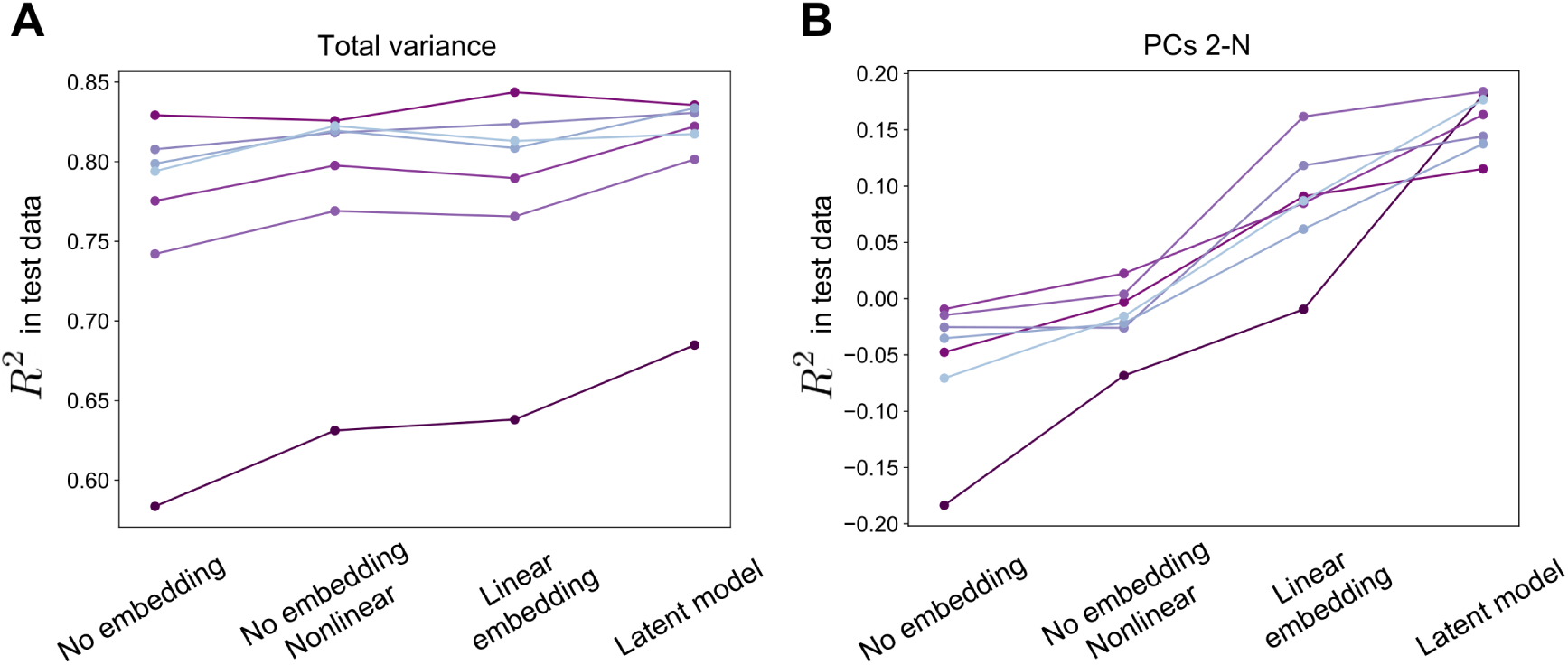
Variance explained in test data for four model categories: “No embedding” (linear pupil regressor), “No embedding, Nonlinear” (nonlinear mapping from pupil regressor), “Linear embedding” (linear mapping from delay embedded pupil), and “Latent model” (nonlinear mapping from delay embedded pupil). Left, total variance explained; right, variance explained along PCs 2-N. Compare also with simulation in Fig. S3 C&E.

**Figure S5:**
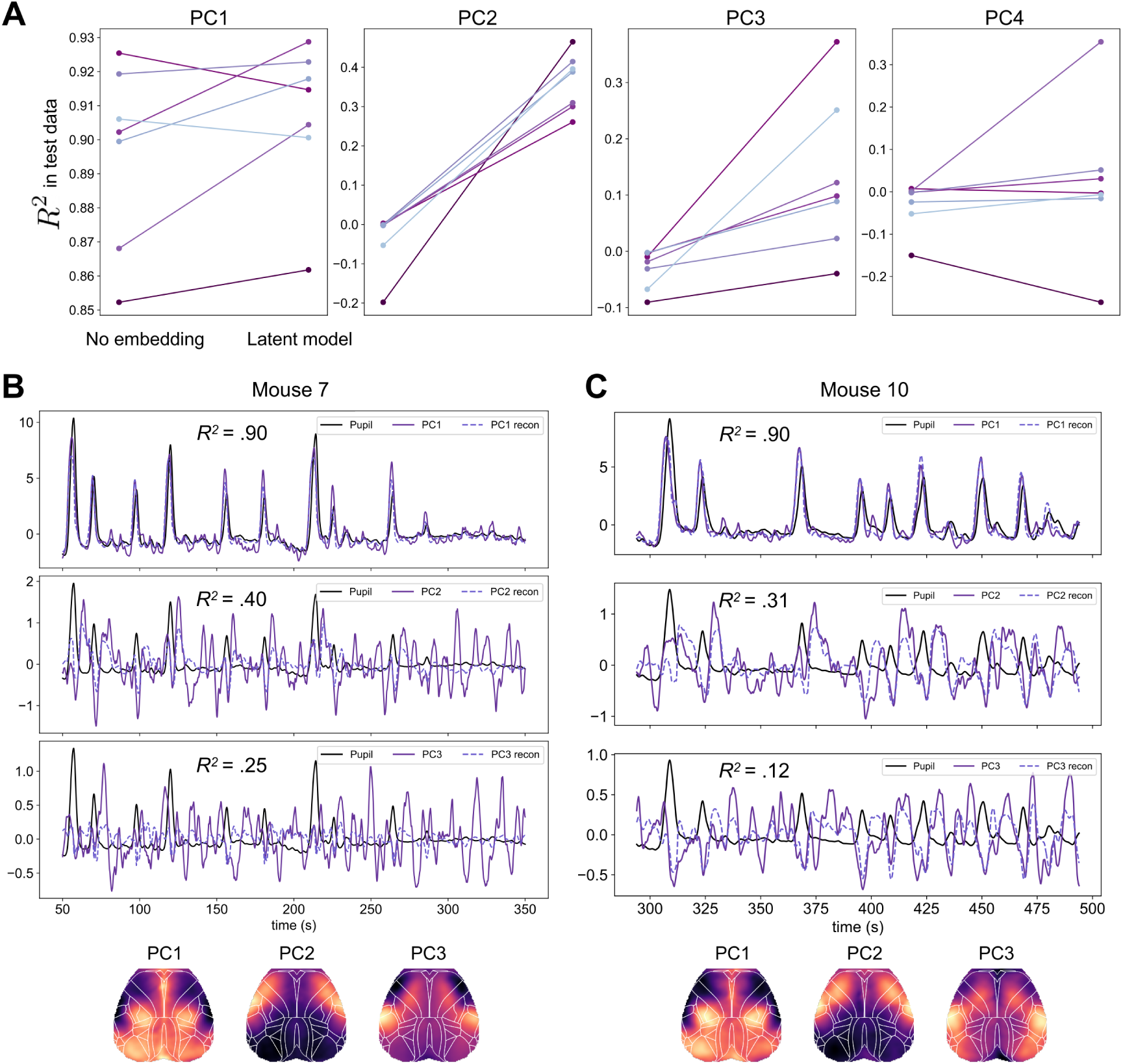
Pupil delay embeddings explain multidimensional variance in widefield calcium images. **A** Variance explained along the dimensions spanned by each of the first four principal components from each mouse. **B-C** Principal component (PC) time series and their prediction from pupil delay embeddings in test data, shown for two example mice. PC spatial maps corresponding to each time series are shown underneath. Higher-order PCs (here, 2 and 3) retain clear temporal relationships to PC1, enabling some of this multidimensional variance to be captured from a time-invariant mapping from the pupil-derived latent space.

**Figure S6:**
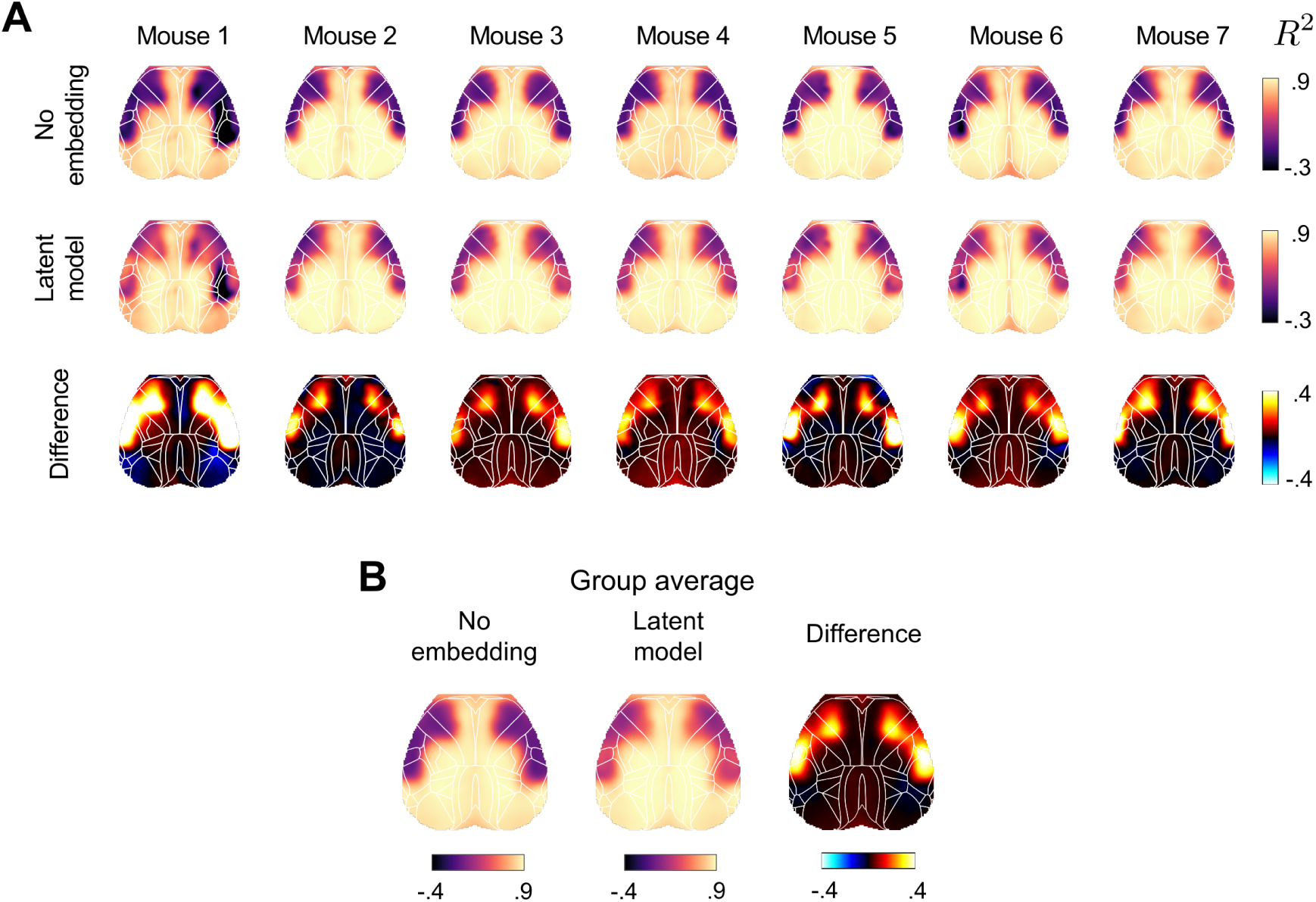
*R*^2^ maps in held-out data corresponding to the “No embedding” and “Latent model” predictions, shown for each mouse (A) and averaged across mice (B). Dynamical information consistently improved explanatory power, particularly within a bilateral set of premotor cortical regions. This pattern recalls prior results in humans (cf. [117, 165]).

**Figure S7:**
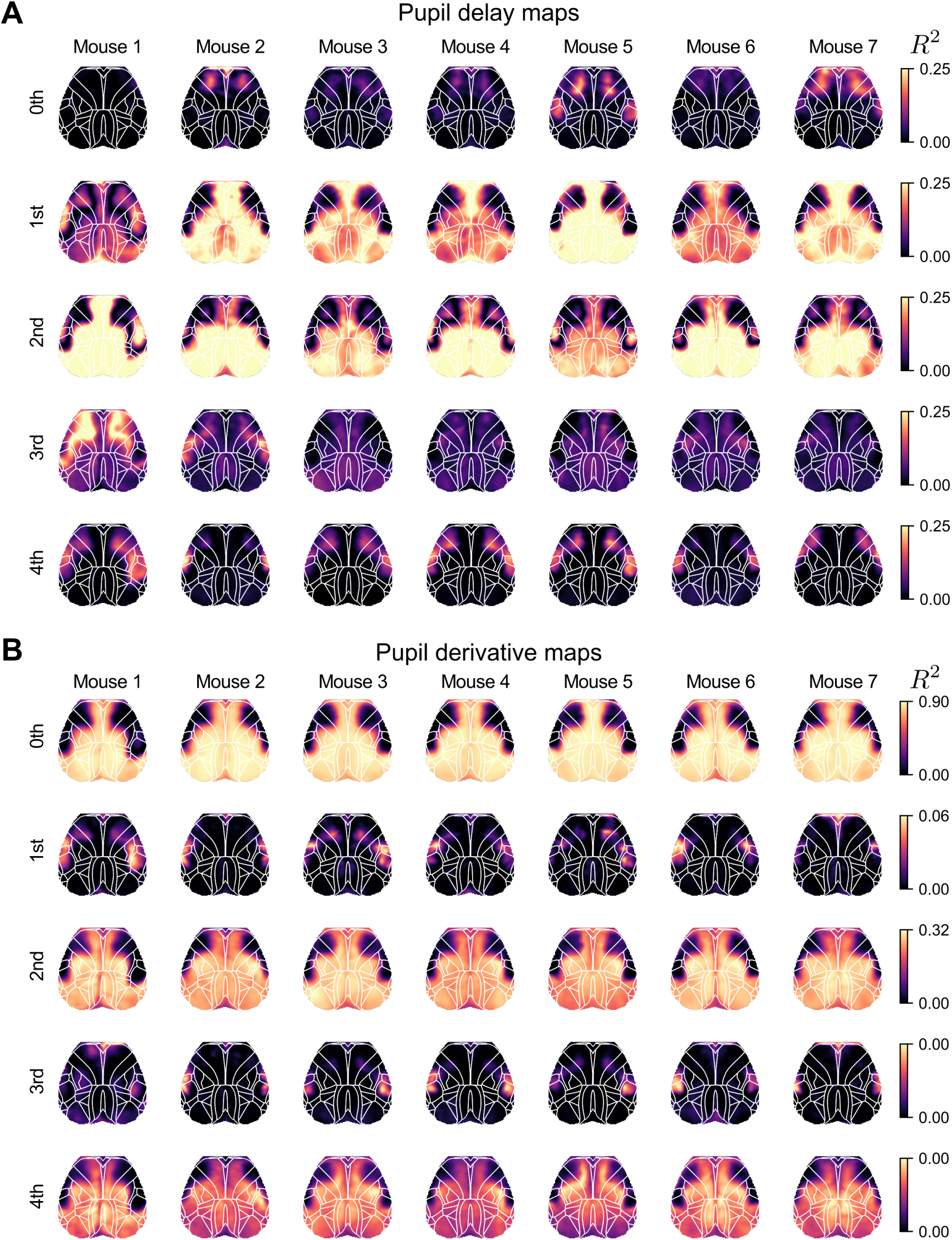
*R*^2^ maps for widefield calcium images as linearly modeled by dynamical features of pupil size (no cross-validation). **A** *R*^2^ maps derived from the pupil delay embedding (i.e., the Hankel matrix) projected onto successive Legendre polynomials. Projections onto Legendre polynomials (i.e., the Legendre coordinates 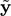 from Eq. (5)) essentially define convolutions of pupil diameter with increasingly high-frequency information from the recent past. The 0th-order polynomial effectively captures the amplitude envelope of pupil diameter (cf. widefield maps obtained from cholinergic indicators [20], which also correlate with pupil amplitude envelope [94]). **B** *R*^2^ maps derived from successive temporal derivatives of pupil diameter (0th order derivative corresponds to the original pupil diameter time series).

**Figure S8:**
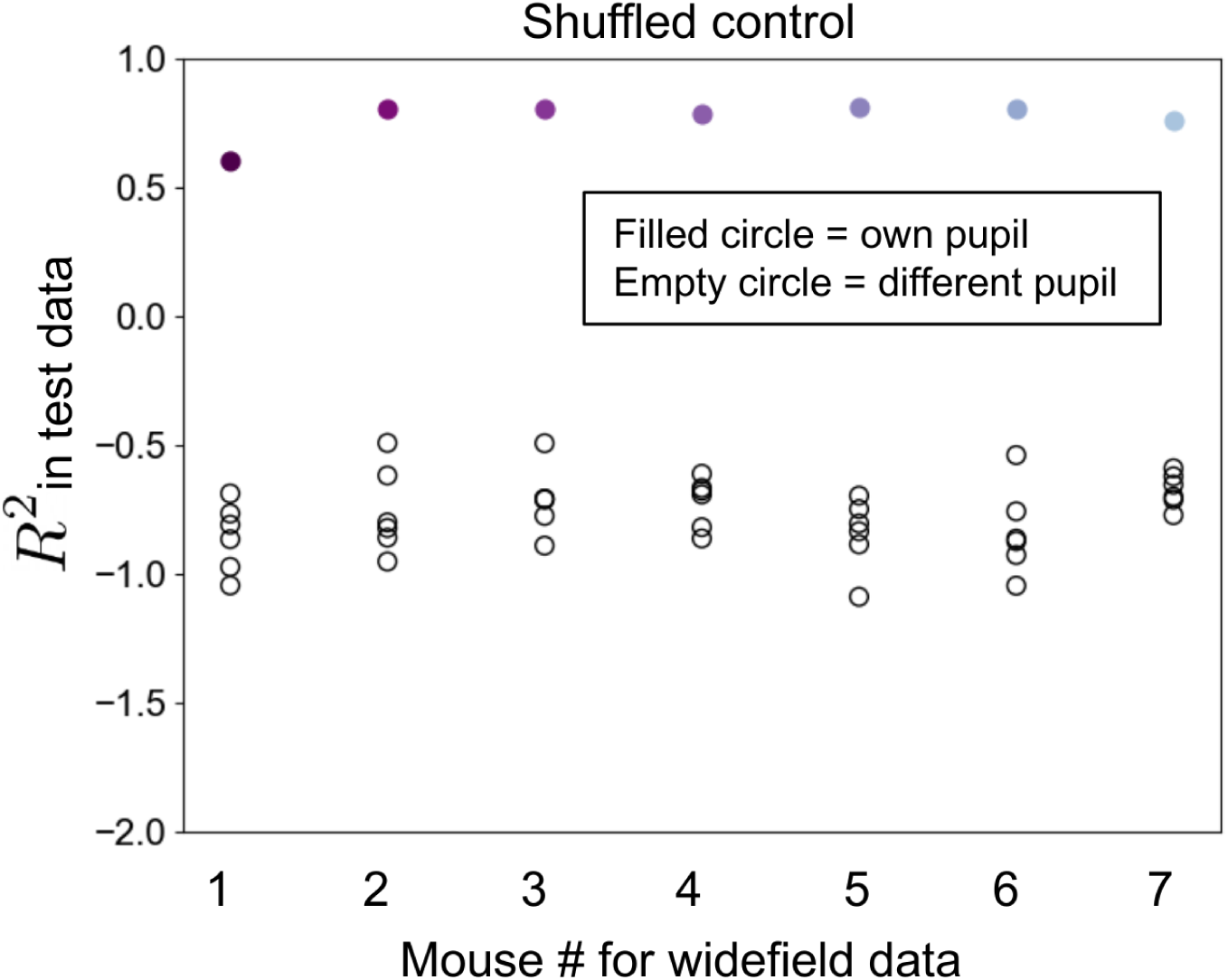
Shuffled control analysis for prediction of widefield calcium data from pupil dynamics (compare with Fig. 2A). Models were trained according to the standard leave-one-out pipeline. However, at test time, widefield data from the held-out mouse was predicted from not only the held-out mouse’s own pupil data (filled circles), but also, separately, from the pupil data from each of the other mice (empty circles). For all mice, explained variance is high when using the mouse’s own pupil timecourse (filled circles). In contrast, explained variance is invariably poor when attempting to predict widefield data from pupil dynamics obtained from another mouse (empty circles; negative *R*^2^ values occur when the prediction performs worse than simply predicting the mean value of each pixel). This result clarifies that good performance in held-out data is not trivially obtained with any complex model.

**Figure S9:**
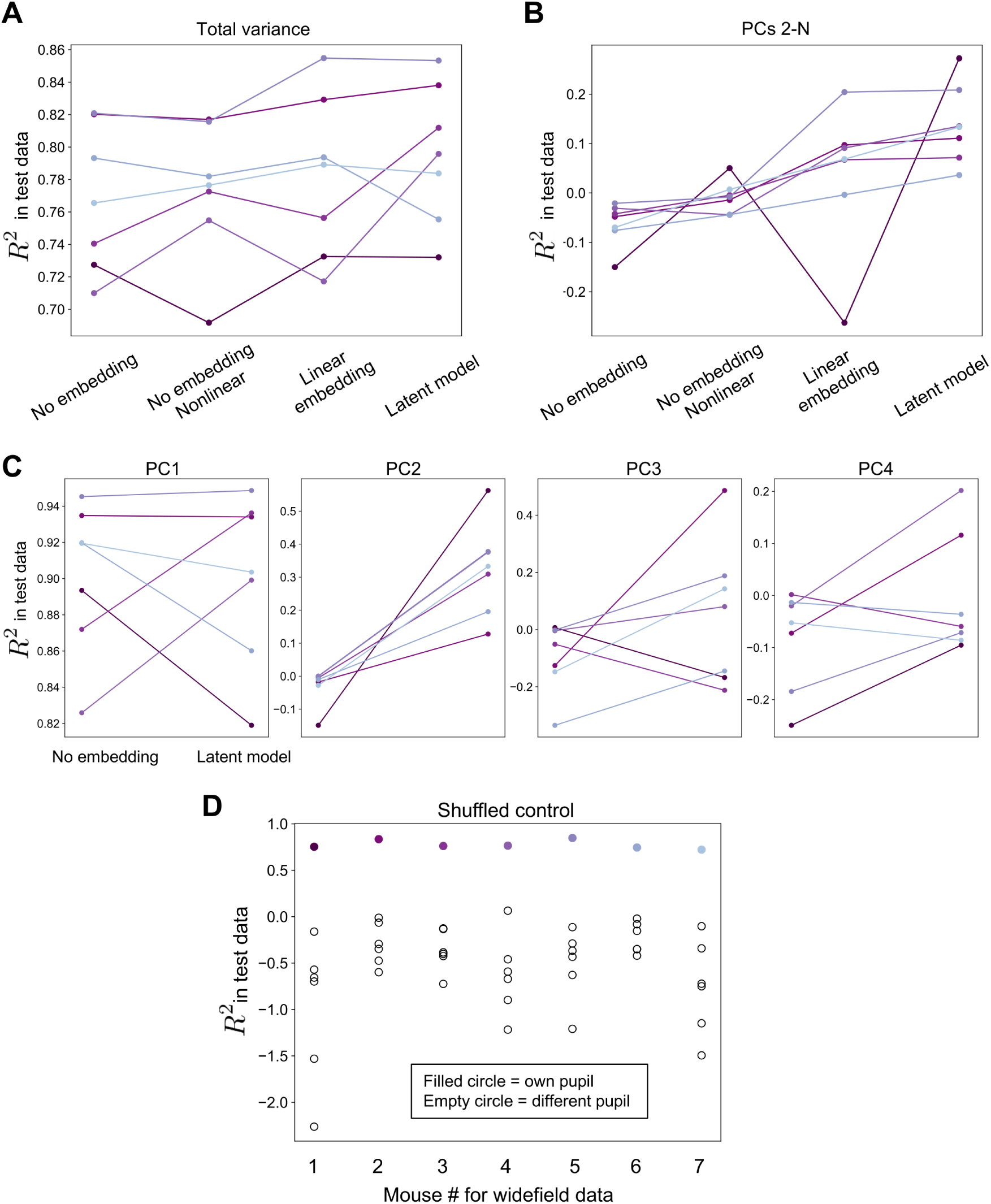
Results from the within-mouse model pipeline. Each mouse’s dataset was split into train and test sets, as detailed in the Methods. **A-B** Same as in Fig. S4. **C** Same as in Fig. S5A. **D** Shuffled control analysis, as in Fig. S6, but adapted for the within-mouse model pipeline. Accordingly, for each mouse’s test data, we trained seven models to predict the widefield data from pupil dynamics, with each model using the pupil time series from the training set of a different mouse. Explained variance (*R*^2^) is reported for each model in test data. Filled circles correspond to models trained on the test mouse’s own training data, as in A-C.

**Figure S10:**
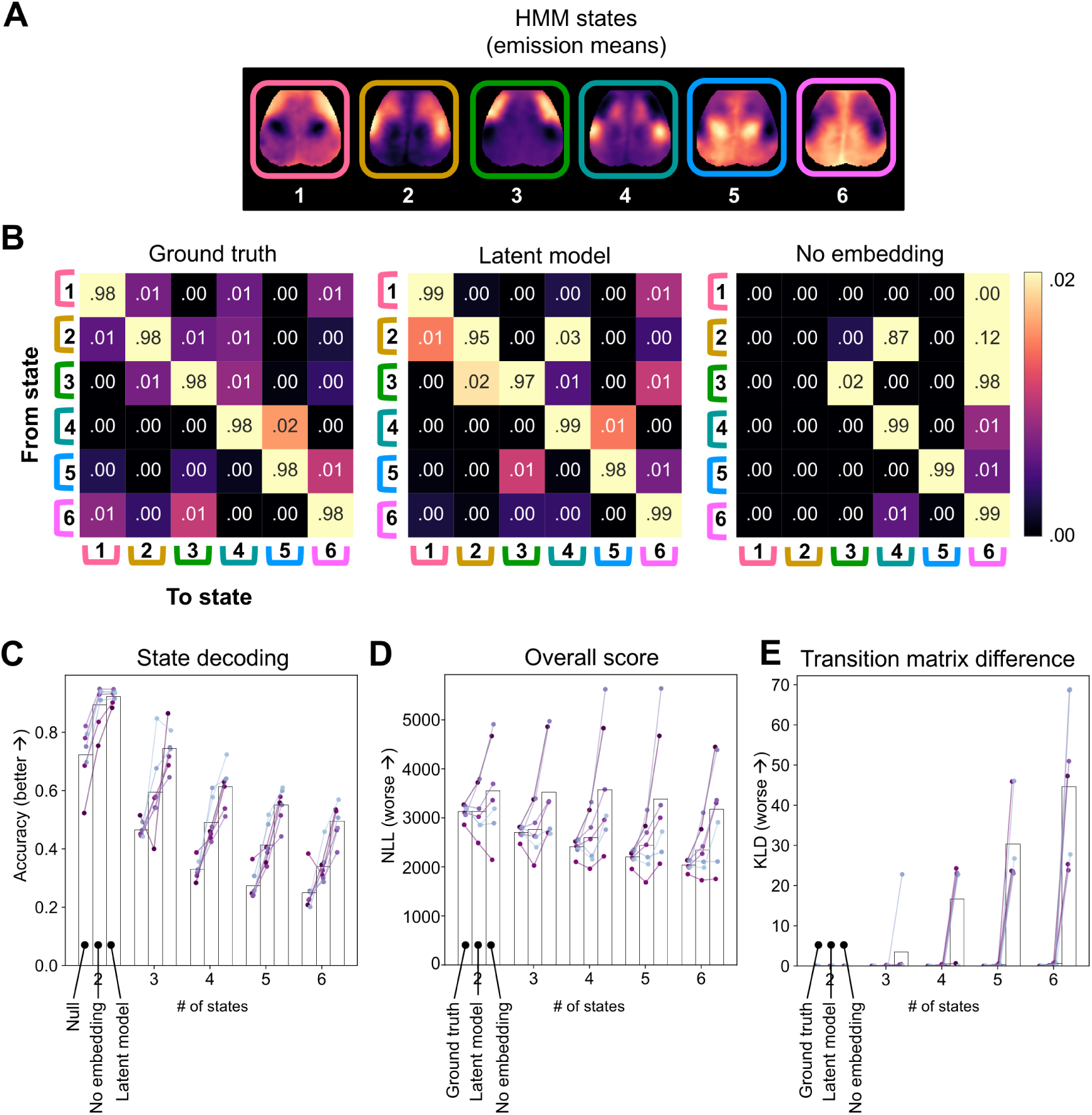
Hidden Markov models (HMMs) reveal that the latent model recapitulates spatially structured dynamics that are not captured by instantaneous pupil diameter. **A** Six HMM “states” for a sample mouse (same as in Fig. 2D), computed as the mean of the Gaussian emission associated with each hidden state. **B** HMM transition matrices involving the HMM states in A for the sample mouse, computed from the original data (left), the data reconstructed through the Latent model (middle), and the data reconstructed through the No embedding model (right). See Methods for model fitting details. Note high probabilities along the superdiagonal (elements immediately to the right of the principal diagonal) indicating a canonical cycle among states in this mouse (cf. Fig. 2D). This structure is less apparent in the No embedding reconstruction (right). **C-E** Evaluation metrics computed over a range of latent states (2-6) for all seven mice. **C** Decoding accuracy, as in Fig. 2E. Null predictions are based on repeatedly choosing the most common state in the ground truth HMM for each time point. Greater accuracy indicates greater agreement of latent state labels with labels obtained from the original HMM. **D** Total negative log-likelihood (NLL) of the ground truth or reconstructed datasets based on the HMM fit to the original data. Greater NLL indicates data are less probable according to ground truth generative model. **E** KL divergence of the ground truth and reconstructed transition matrices from the original HMM transition matrix. Greater KL divergence indicates greater difference from the ground truth transition matrix.

**Figure S11:**
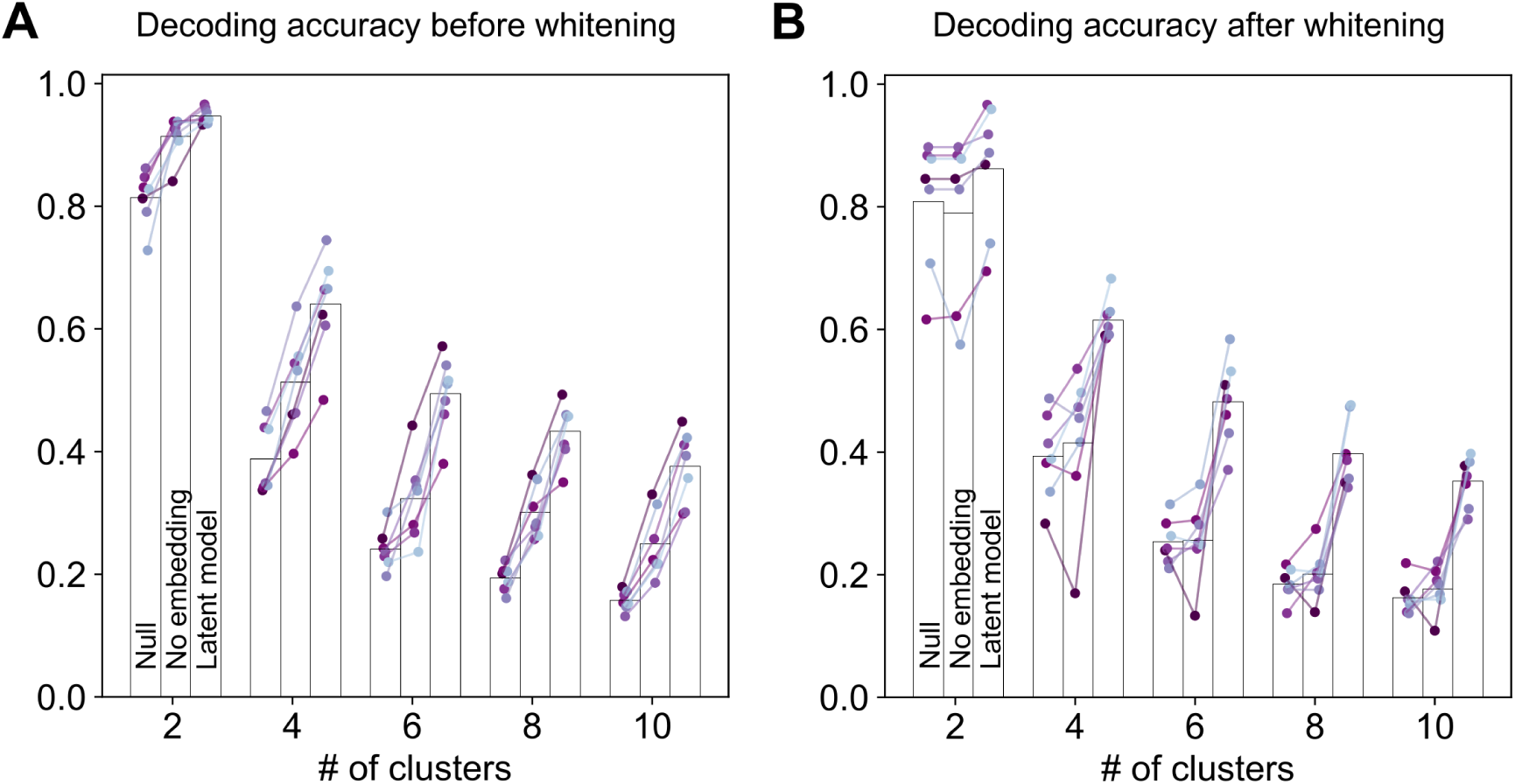
Decoding “brain states” from pupil dynamics. Comparison of decoder accuracy in properly identifying GMM cluster assignments for each image frame based on the most frequent assignment in the training set (“Null” decoder), the “No embedding” model, or the “Latent model”. **A** Same as in Fig. 2E, but for up to 10 clusters and using a spherical rather than full covariance prior (Methods). **B** Same as in (A) but after whitening transformation (i.e., after projection onto the top 3 PCs, each dimension was set to unit variance). The whitening transformation highlights the ability of the delay embedding to capture genuinely multidimensional information, which is otherwise masked by the dominant PC1.

**Figure S12:**
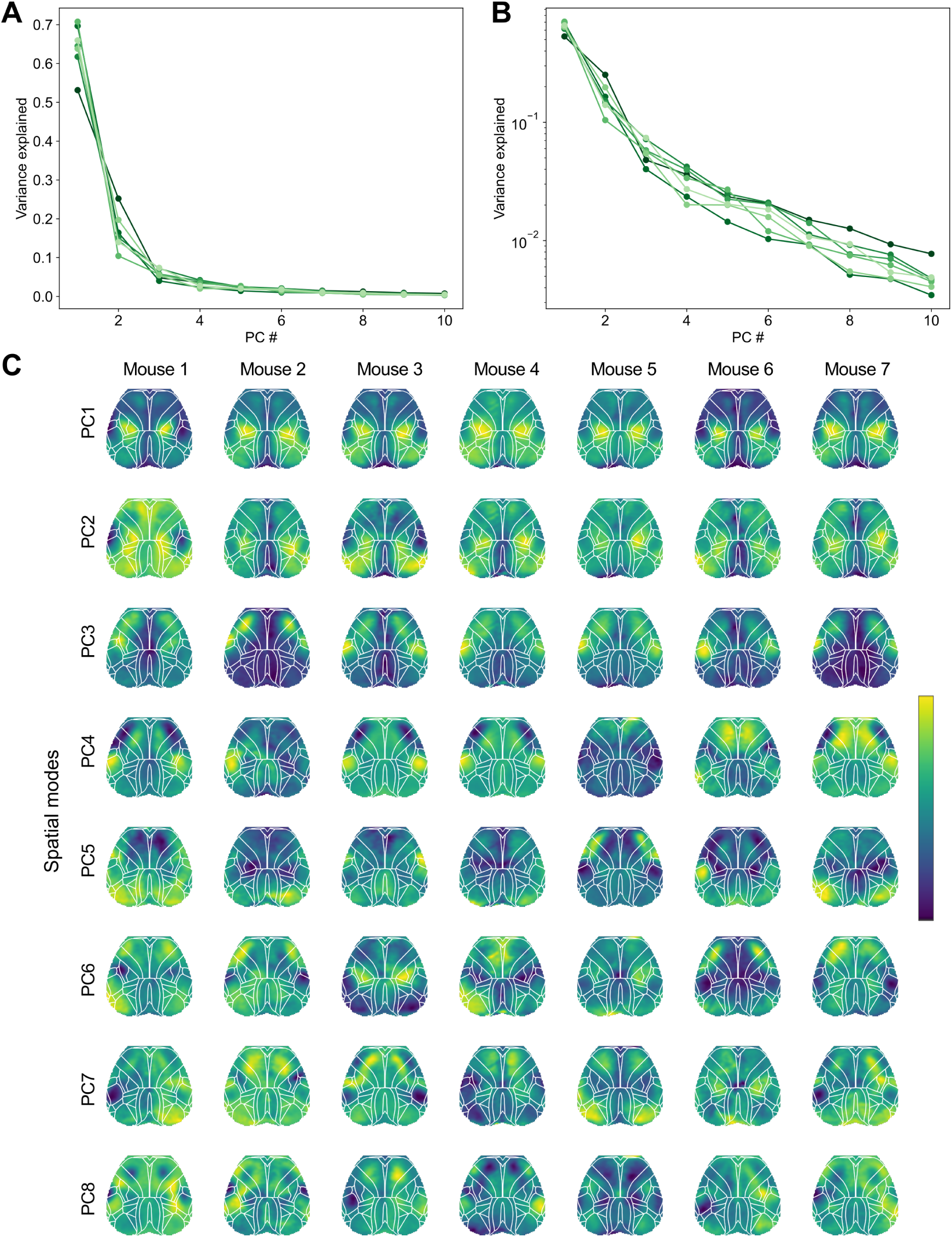
SVD results for widefield FAF images (as in Fig. S1). **A-B** Scree plots for each of the seven mice, displayed on a linear (A) or log-transformed (B) y-axis. For all mice, images were dominated by the first principal component. **C** Spatial topographies corresponding to the top 8 PCs (arbitrary units). As the sign of each PC is arbitrary, some PCs were sign-flipped to improve visual correspondence across mice.

**Figure S13:**
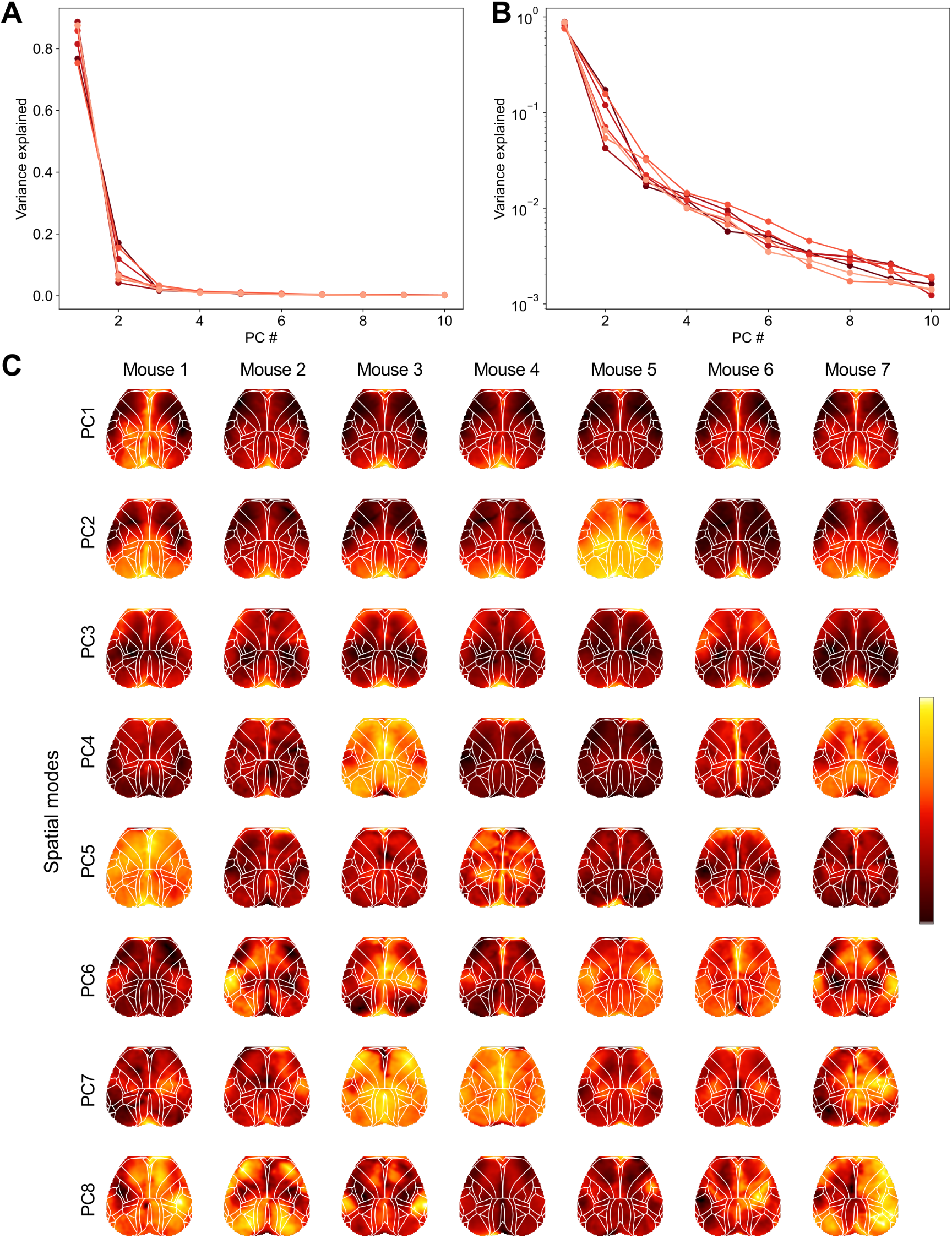
SVD results for widefield hemoglobin images (as in Figs. S1 & S12). **A-B** Scree plots for each of the seven mice, displayed on a linear (A) or log-transformed (B) y-axis. For all mice, images were dominated by the first principal component. **C** Spatial topographies corresponding to the top 8 PCs (arbitrary units). As the sign of each PC is arbitrary, some PCs were sign-flipped to improve visual correspondence across mice.

**Figure S14:**
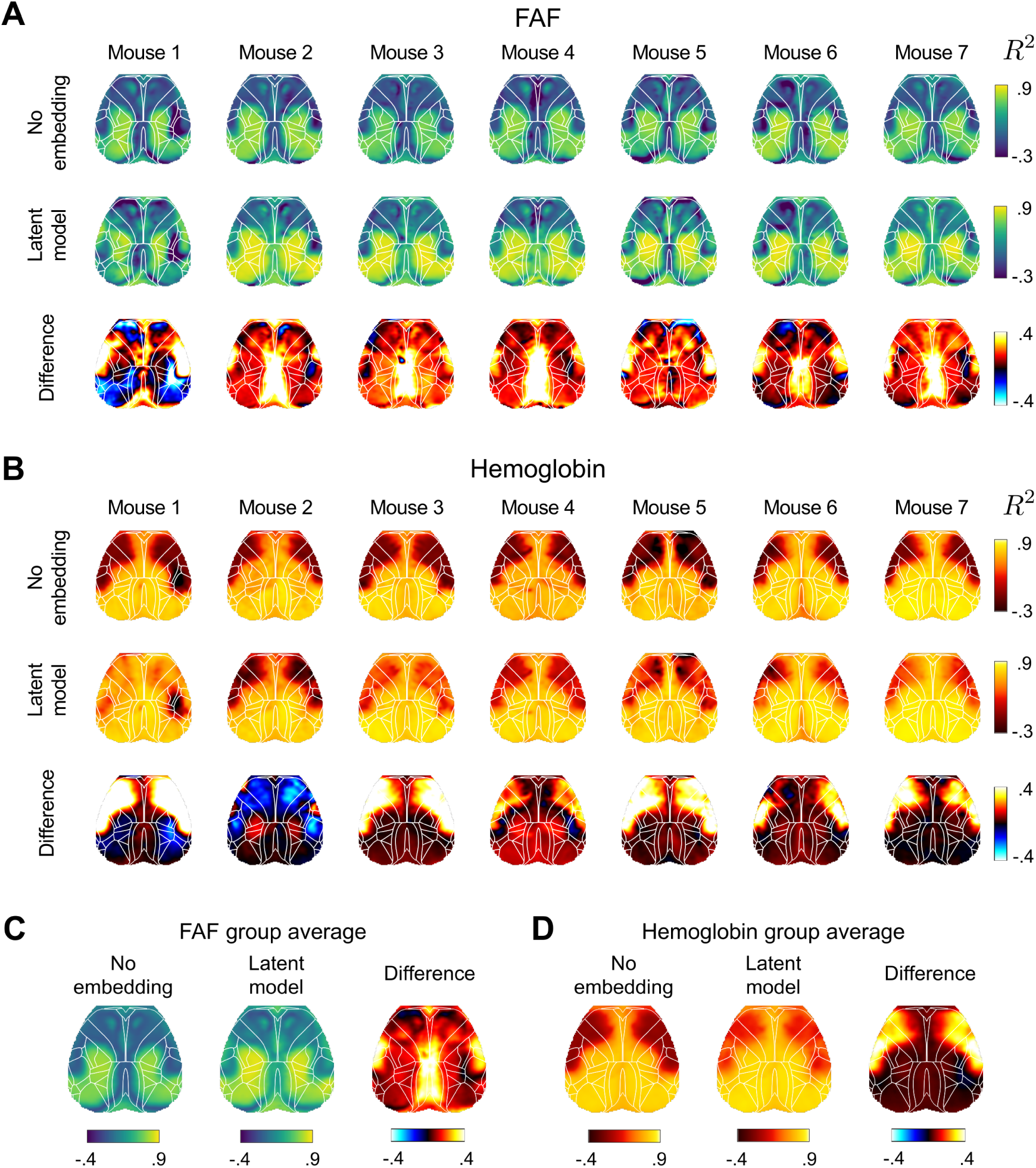
*R*^2^ maps in held-out data corresponding to the “No embedding” and “Latent model” predictions, shown for FAF (A, C) and hemoglobin (B, D), for each mouse (A, B) and averaged across mice (C, D). See Fig. S6 for calcium results.

**Figure S15:**
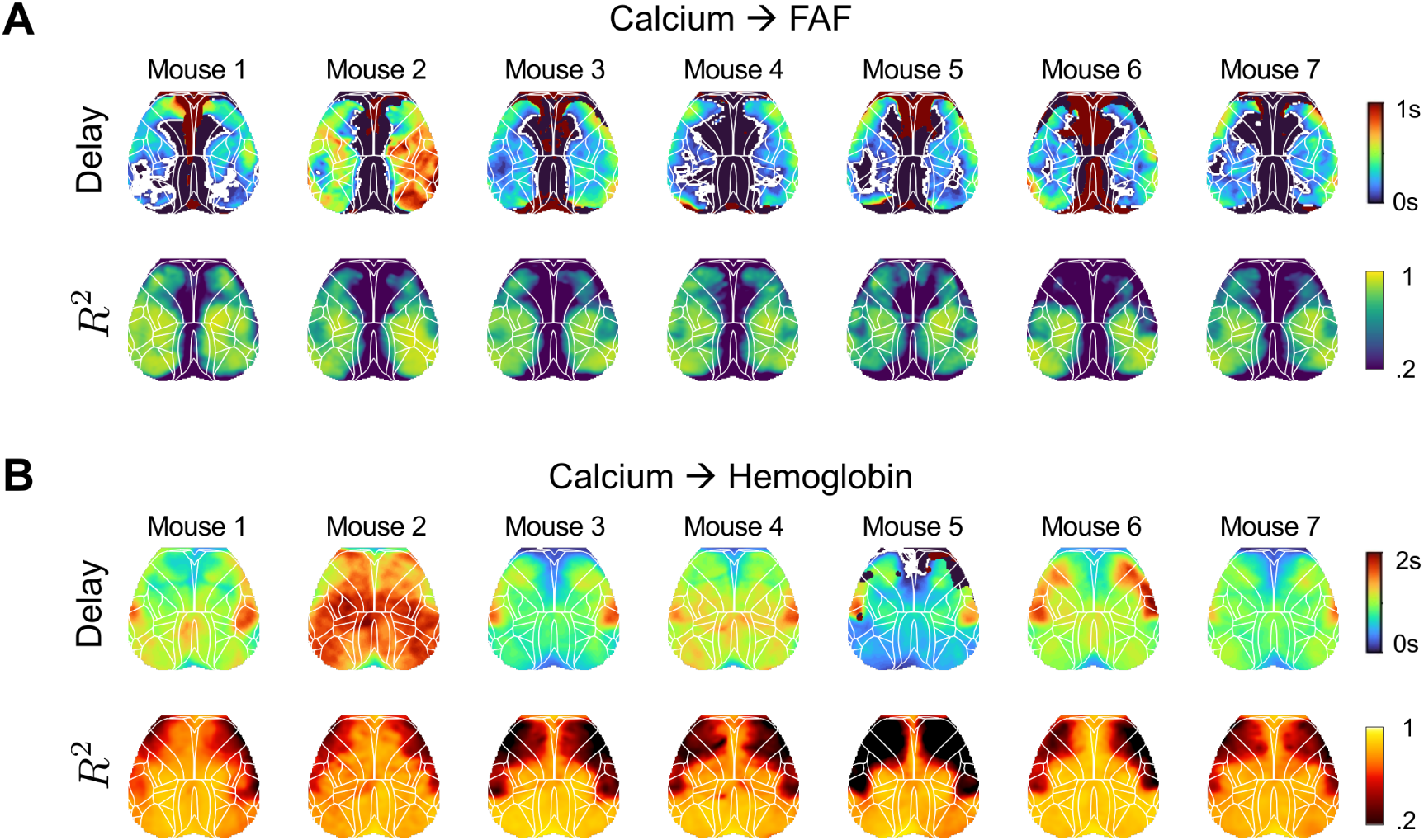
*R*^2^ maps for widefield FAF (A) and hemoglobin (B) recordings linearly modeled at each pixel according to the widefield calcium measurements recorded at the same pixel. *R*^2^ values (lower panels) are reported at the time shift (upper panels) optimizing the cross-correlation function between the calcium and FAF (or hemodynamic) recording at that pixel. Note that these model results are reported as is, i.e. without cross-validation. Extremal or missing values in the delay maps are generally due to the absence of a single, well-defined peak in the cross-correlation function within the allowable range (±5 sec). This is commonly caused by biphasic patterns in the cross-correlation function. Note that delay embedding avoids this issue by combining multiple lagged copies of the predictor variable into a single, multidimensional predictor variable, rather than requiring a single lag-optimized predictor variable.

**Figure S16:**
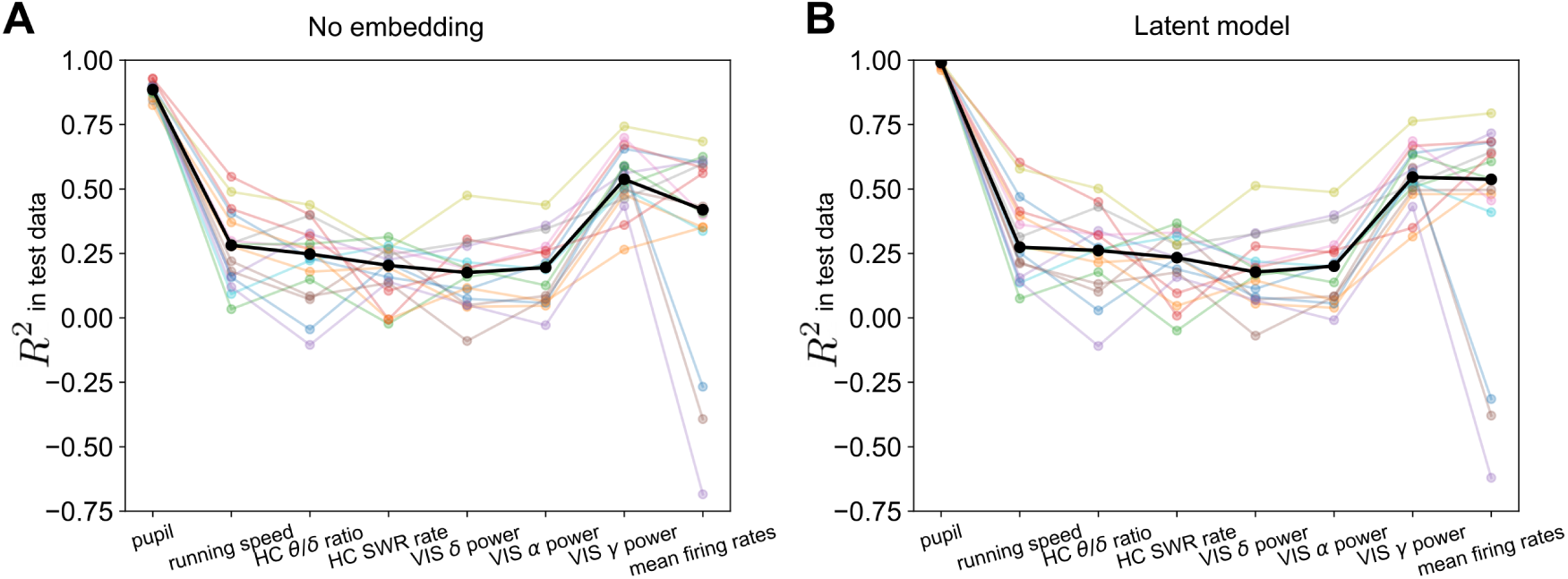
Leave-one-out cross validation analysis in Allen Brain Observatory data. Colored lines correspond to individual mice; black lines reflect median values across mice.

